# Evolutionary games with environmental feedbacks

**DOI:** 10.1101/493023

**Authors:** Andrew R. Tilman, Joshua Plotkin, Erol Akçay

## Abstract

Strategic interactions form the basis for evolutionary game theory and often occur in dynamic environments. The various strategies employed in a population may alter the quality or state of the environment, which may in turn feedback to change the incentive structure of strategic interactions. This type of feedback is common in social-ecological systems, evolutionary-ecological systems, and even psychological-economic systems – where the state of the environment alters the dynamics of competing types, and vice versa. Here we develop a framework of “eco-evolutionary game theory” that permits the study of joint strategic and environmental dynamics, with feedbacks. We consider environments governed either by a renewable resource (e.g. common-pool harvesting) or a decaying resource (e.g. pollution byproducts). We show that the dynamics of strategies and the environment depend, crucially, on the incentives for individuals to lead or follow behavioral changes, and on the relative speed of environmental versus strategic change. Our analysis unites dynamical phenomena that occur in settings as diverse as human decision-making, plant nutrient acquisition, and resource harvesting. We discuss the implication of our results for fields ranging from ecology to economics.

---

In many interactions, an individual’s payoff depends on both her own strategy or type, as well as the strategic composition in the entire population. Such interactions arise across a range of disciplines, from micro-economics to animal behavior, and they have been analyzed using game theory (Nash, 1950; Maynard Smith, 1982). Game-theoretic analysis of competing types typically assumes that the form of strategic interaction is fixed in time, or that it depends on an independent exogenous environment. Real-world systems, however, often feature bi-directional feedbacks between the environment and the nature of strategic interactions: an individual’s payoff depends not only her actions relative to the population, but also on the state of the environment, and the state of the environment is influenced by the actions adopted by all individuals in the population.

Reciprocal feedbacks between strategic interactions and the environment play out in many complex systems with broad biological and societal relevance (Sethi and Somanathan, 1996; Tavoni et al., 2012; Levin et al., 2013; Tilman et al., 2018; Mullon et al., 2018; Estrela et al., 2018). In fisheries, for example, the relative reward of a high-intensity versus a low-intensity harvesting strategy depends upon the current biomass of the fish stock; and, conversely, stock dynamics depend on the frequencies of these two harvesting strategies(Tilman et al., 2017). In ecology, species strategies and interactions determine their competitive balance whilst also changing the abiotic environment and, in turn, the nature of competition. For example, symbiotic nitrogen fixation in plants increases local nitrogen availability over time, altering the competitive environment (Menge et al., 2008). Conversely, nutrient availability will change incentives for nutrient exchange in such symbioses (Grman et al., 2012). Likewise, global climate change is caused by the strategic decisions of individuals, corporations, and nations, which have long-term environmental repercussions that will, in turn, alter the strategic landscape those parties face. Feedbacks even occur in psychology, where deliberative decision-making can beneficially shape the shared environment of social norms and institutions, but that environment then sets the stage for the success of less costly, non-deliberative decision-making (Rand et al., 2017). Examples from these diverse fields share the common feature that feedbacks between strategies and the environment fundamentally alter dynamical predictions (Peck and Feldman, 1986; Worden and Levin, 2007; Cortez and Ellner, 2010; Akçay and Roughgarden, 2011; Huang et al., 2012; Stewart and Plotkin, 2014; Weitz et al., 2016; Akçay, 2018; Cortez et al., 2018; Hilbe et al., 2018; Patel et al., 2018; Shao et al., 2019). Understanding such systems requires a framework for studying game-theoretic dynamics when the nature of competitive interactions influences the environment, and conversely.

Here we develop a general framework for eco-evolutionary games. We analyze the case of a two-strategy game linked to a renewable or decaying resource. We derive stability criteria for all strategic/environmental equilibria, and we derive the conditions that permit cyclic dynamics, bistability, or a stable equilibrium that supports multiple co-existing types. We show that environmental feedbacks alter equilibrium outcomes and expand the suite of dynamical possibilities. Importantly, we can characterize the large range of possible dynamical behavior in terms of a few easily interpretable quantities: incentives to lead or follow strategic changes in the population. We find that cyclical dynamics arise in a restricted subset of game structures, and that in these regions, the dynamical behavior depends critically on the relative timescale of strategic versus environmental change.

## 1 Eco-evolutionary games

Eco-evolutionary games occur when evolutionary game dynamics are environmentally coupled. First, consider a set *S* of different strategies that can be employed in a system of interest. Depending on the system being studied, the strategies may be defined as a set of alternative resource extraction technologies, or various nutrient acquisition strategies of plants, or different physiologies or morphologies of organisms, or different cognitive types that excel in different environments. We assume that an individual’s fitness from adopting strategy *s*_*i*_ ∈ *S* is *π*_*i*_(**x**, *n*). The individual’s fitness is a function of her strategy, the frequency of all *K* strategies in the population, **x** = [*x*_1_, *x*_2_, …, *x*_*K*_], which we call the population strategy profile, and also the state of the environment, *n*. Depending upon the context, the environmental state might correspond to the concentration of a vital nutrient, the biomass of species in a community, the concentration of greenhouse gasses (or other pollutants) in the atmosphere, or the quality of social norms. Since the fitness of an individual depends on the strategies employed by other individuals, the setting we have described is game-theoretic in nature.

We study the dynamics of the frequencies of strategies with the replicator equation (Taylor and Jonker, 1978), writing the rate of change of the frequency of strategy *s*_*i*_ as

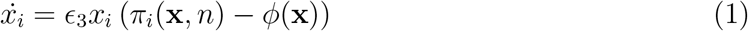

where 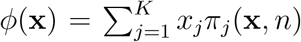 is the mean fitness of the population and ∊_3_ is a parameter that describes the speed of strategy dynamics. This equation implies that the frequency of a strategy increases when the fitness of those who adopt it is greater than the average fitness of the population.

We have described a game-theoretic interaction that is environmentally dependent. But we are interested in systems which, furthermore, contain environmental feedbacks. Such feedbacks arise when strategies have an impact on the environment. The impact on the environment is channeled through a function, *h*(**x***, n*), which aggregates the influence of the current population strategy profile on changes to the environmental variable *n*. The environmental factor *n* may also have its own intrinsic dynamic governed by *f* (*n*), which describes the intrinsic rate of change of the environmental variable as a function of the current environmental state. Depending on the system of study, these intrinsic dynamics could describe food webs (if modeling a higher-dimensional environmental state), soil weathering, or earth systems processes. This results in environmental change governed by

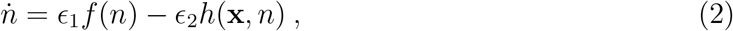

where ∊_1_ and ∊_2_ determine the speeds of the intrinsic environmental dynamics (independent of strategies played) and of the extrinsic impact of strategies on the environment, respectively. In total, this describes a system where evolutionary dynamics are reciprocally linked with the environment in a dynamic eco-evolutionary game. The model is described by a system of *K* differential equations (1 environmental equation and *K −* 1 strategy equations, since **x** lies on a simplex).

For a two-strategy game with an environmental feedback we can write the eco-evolutionary system as

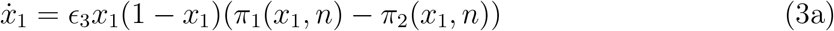

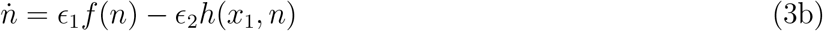

because in this case *x*_1_ = 1 *− x*_2_. This framework features three different timescales: the timescale of intrinsic dynamics of the environment, ∊_1_, the timescale of the environmental impact of the strategies currently employed in the population, ∊_2_, and the timescale of strategy update dynamics (strategy evolution) in the population, ∊_3_. We can normalize the first two timescales relative to the third so that we drop ∊_3_, without loss of generality. This framework allows for non-linearity in the payoff structures as well as in environmental impact and intrinsic dynamics, so that the space of models and potential dynamics is vast. We first focus on the subset of models in which payoffs are linear in the state of the environment and the strategy frequencies, and we later consider non-linear payoffs.

## 2 Linear eco-evolutionary games

In this section we describe a class of two-strategy eco-evolutionary games where the payoffs to individuals are linear in both the population strategy profile, *x*, and in the environmental state, *n*. We will show that several important models from disparate fields are instances of linear eco-evolutionary games, in which we can write the payoffs in terms of a matrix

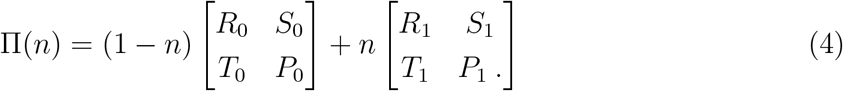

Here the state of the environment, *n*, is normalized to fall between 0 and 1, and the entries of the two matrices correspond to the payoffs of the game under rich (*n* = 1) and poor (*n* = 0) environmental states. Using Π(*n*), we write the payoffs for using strategy 1 and strategy 2 as *π*_1_(*x, n*) and *π*_2_(*x, n*), respectively, where *x* denotes the fraction of the population that plays strategy 1. We systematically analyze games of this form coupled to environments (resources) with either renewing or decaying intrinsic dynamics.

### 2.1 Eco-evolutionary games with intrinsic resource dynamics

Intrinsic resource dynamics can take many forms, but are broadly categorized as renewing or decaying. We consider two different environmental dynamics: (i) a renewable resource where each strategy exerts degradation (or harvesting) pressure on the resource stock, and (ii) a decaying resource that is produced as a by-product of each strategy.

We first suppose there is a resource stock, *m*, that grows logistically in the absence of consumption or harvesting, and is diminished by harvesting or consumption associated with the strategies in a game. Let *e*_*L*_ and *e*_*H*_ be the resource harvest effort of strategies Low and High, respectively, with *e*_*L*_ < *e*_*H*_. The resource dynamics are then governed by

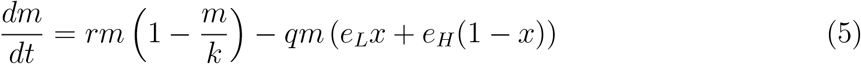

where *x* is the fraction of the population playing strategy L, *r* is the intrinsic rate of resource growth, and *k* is the resource carrying capacity. Here *q* is a parameter that maps resource degradation pressures (or harvesting efforts) (*e*_*L*_, *e*_*H*_) into the rate of reduction in the resource. We assume that environmental impact rates are restricted so that *m* will be positive at equilibrium. This implies that *e*_*H*_ ∈ (0, *r/q*) and *e*_*L*_ ∈ [0, *e*_*H*_), which spans the effort the leads to maximum sustainable yield as well as the bio-economic effort level associated with open access and the tragedy of the commons. And so such models are well suited to address questions of environmental stewardship or over-exploitation.

The resource stock *m* can assume any non-negative value. Regardless of its initial value, though, *m* will eventually fall between its equilibrium values when either high- and low-effort strategies dominate the population. Therefore, we can transform *m* to an environmental state *n* in our framework, which is bounded between 0 and 1, using a simple linear transformation between these two extreme equilibrial values (See supplementary information section S1.1).

Now, using the payoff matrix from equation 4 we can write our eco-evolutionary system as

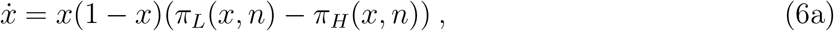

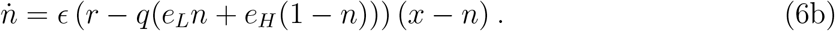

In terms of our general framework (equation 2) we have *f* (*n*) = *−* (*r − q*(*e*_*L*_*n* + *e*_*H*_ (1 *− n*))) *n* and *h*(*x, n*) = *−* (*r − q*(*e*_*L*_*n* + *e*_*H*_ (1 *− n*))) *x*. Here we have assumed that the timescales of intrinsic resource dynamics and resource harvesting are the same (i.e., ∊_1_ = ∊_2_ = ∊), so that we have only one relative timescale remaining. In other words, ∊ quantifies the rate of environmental dynamics (both intrinsic and extrinsic) compared to the rate of strategy dynamics.

We can make a similar mapping for a model with a decaying resource, such as a pollutant. Let *m* denote the pollutant level in the environment. Each strategy played in the evolutionary game produces the pollutant as a byproduct at some rate. Let *e*_*L*_ and *e*_*H*_ be the emissions rates of the low emissions and high emissions strategies, respectively. Then we can model the stock of *m* with the differential equation

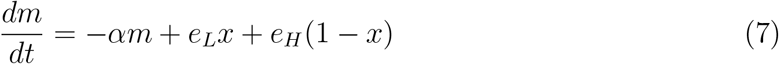

where *x* is the fraction of the population that plays the strategy with low emissions (strategy L), (1 *− x*) is the fraction the plays the high emission strategy and *α* is the decay rate of the resource stock. We can again define *n*, bounded between 0 and 1, as a linear transformation of *m* (see supplementary information section S2.1), yielding dynamics governed by

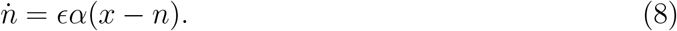

Although decaying versus renewing resources arise in different biological or social contexts, they both yield the same qualitative results and the following analysis of dynamical outcomes applies to both cases.

### Incentives for behavioral change

The dynamics of the eco-evolutionary system above can be understood in terms of four parameter combinations, which have an intuitive interpretations as incentives to change behavior:

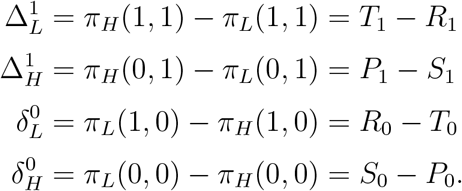

These four parameters will allow us express dynamical outcomes in terms of the incentive to lead or follow strategy change under either a rich or a poor environmental state (Figure 1).

**Figure 1:**
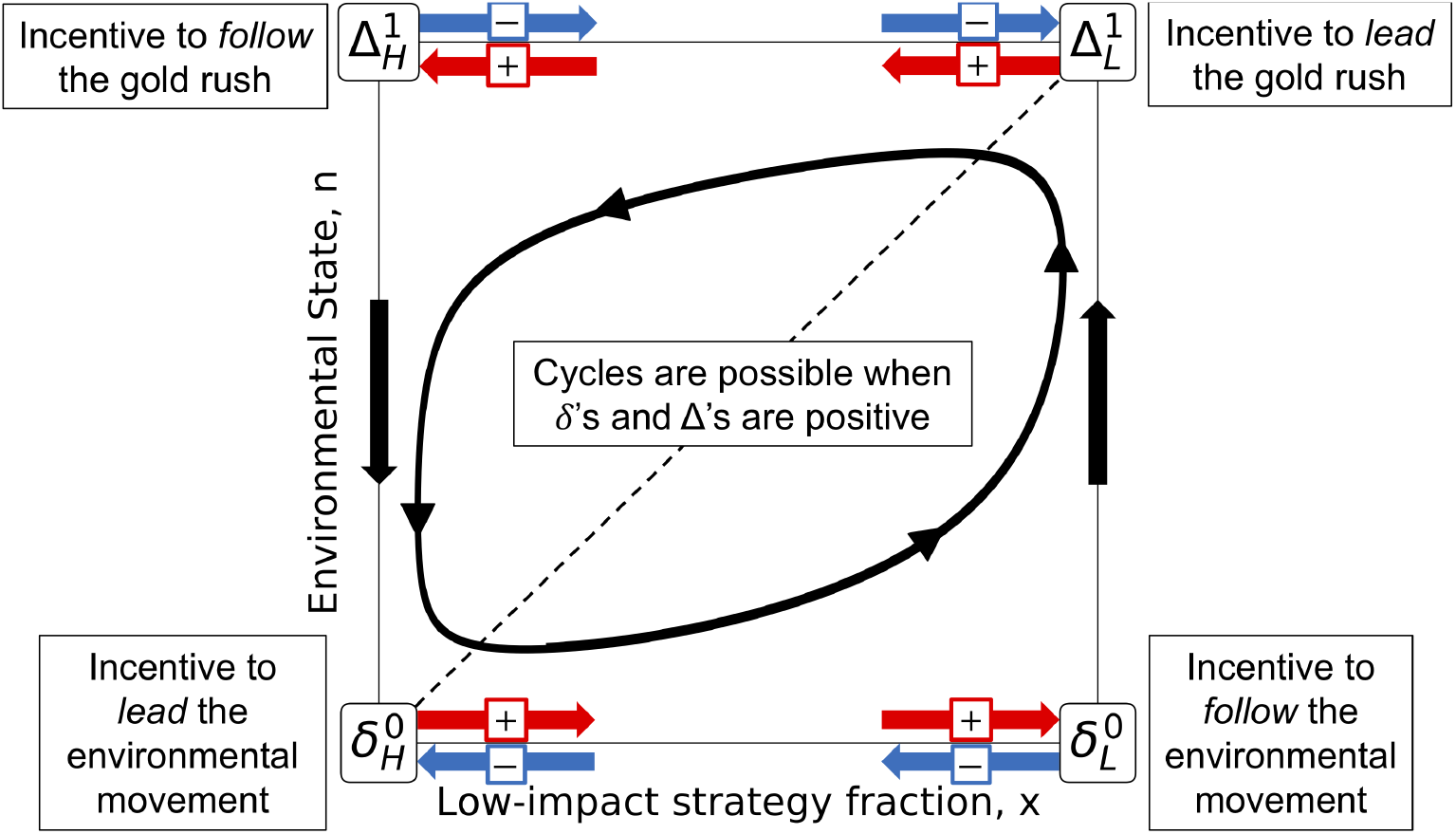
A graphical representation of how incentive parameters in eco-evolutionary games influence dynamics. The horizontal axis of the state space corresponds to the frequency of individuals using the strategy with low impact on the environment, whereas the vertical axis indicates the quality of the environment, *n*, with the dashed line representing the attracting environmental nullcline. Each of the four incentive parameters (*δ*’s and ∆’s) control the direction and magnitude of strategy dynamics at a corner of the state space: strategy dynamics follow the red arrows when the corresponding *δ* or ∆ is positive, and blue arrows when negative. When all are positive, meaning there are incentives to lead and to follow strategic changes, then some form of cyclical dynamics seem plausible. However, we show that all *δ*’s and ∆’s being positive is neither necessary nor sufficient for cyclic dynamics in eco-evolutionary games.

The parameter 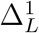 quantifies the incentive to switch to the strategy with high environmental impact (denoted by ∆) given that all other individuals follow the low-impact strategy (denoted by the *L* subscript) and the system is currently in a rich environmental state (denoted by the superscript 1). In the context of socio-ecological systems, 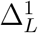 can be interpreted as the incentive to *lead* a "gold rush" – that is, be the first player to switch to a high-impact strategy and reap the rewards of an abundant resource. By contrast, the parameter 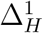 quantifies the incentive to switch to the high-impact strategy under a rich environmental state, given that every other player has already switched. In other words, 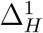 is the incentive to *follow* the gold rush.

The parameter 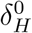 quantifies incentive to switch to the low-impact strategy (denoted by *δ*) when in a poor environmental state and when all other players are following the high-impact strategy. And so we can think of 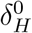 as the incentive to *lead* an “environmental movement" by reducing harvesting of a depleted resource. Finally, the parameter 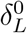 quantifies the incentive to switch to the low-impact strategy given that all other individuals are following the low-impact strategy and the environment is in a poor state. Thus 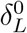 can be seen as the incentive to *follow* the environmental movement.

The verbal descriptions of these four critical parameters apply in the context of a socioecological system, such a fishery. But there are natural alternative interpretations of these parameters for a range of related phenomena across diverse fields.

The general linear evo-evolutionary system has up to four equilibria. There are two equilibria which support a single strategy in the population, at (*x*^∗^, *n*^∗^) ∈ {(0, 0), (1, 1)}; and there are also up to two equilibria that support multiple co-existing strategies in the population, denoted by 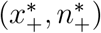 and 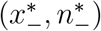. The equilibrium (*x^∗^, n^∗^*) = (0, 0) is stable only if 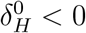, which is intuitively clear from figure 1. Similarly, the equilibrium (*x^∗^, n^∗^*) = (1, 1) is stable only if 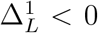. The equilibrium at 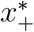 is always a saddle, and thus never stable. Whereas the equilibrium 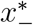 can be either stable or unstable.

We only find persistent cycles in the eco-evolutionary system when the interior equilibrium 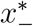 is unstable. Conditions for this equilibrium to be unstable first require

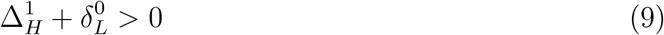

and

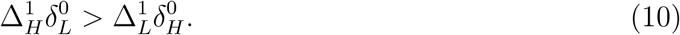

Instability of 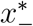 also requires

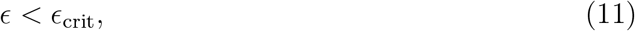

where ∊ is the speed of environmental feedback relative to speed of strategy updating. The value of ∊_crit_ can be expressed analytically in terms of the parameters of the system, and it differs slightly for renewing versus decaying resource feedbacks (See Supplementary information, sections S1.1 and S2.1).

### Dynamical regimes

Strategy-environment dynamics exhibit several different qualitative regimes, depending on the incentives to switch strategies (*δ*’s and ∆’s) and on the timescale separation, ∊, between strategy evolution and environmental impacts.

When there is no incentive to lead either the environmental movement or the gold rush 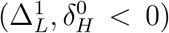, as in Figure 2a, then both edge equilibria are stable, and only the saddle equilibrium falls within the state space. This means that the dynamics in this regime will exhibit bistability – with attraction to a population composed entirely of one or another strategy, depending upon the initial conditions. This result is intuitive because, in this regime, there is no incentive for individuals to be leaders of change in either the low-quality or high-quality environmental state. Therefore the system will eventually be dominated by one or the other strategic type, with the corresponding environmental equilibrium.

**Figure 2:**
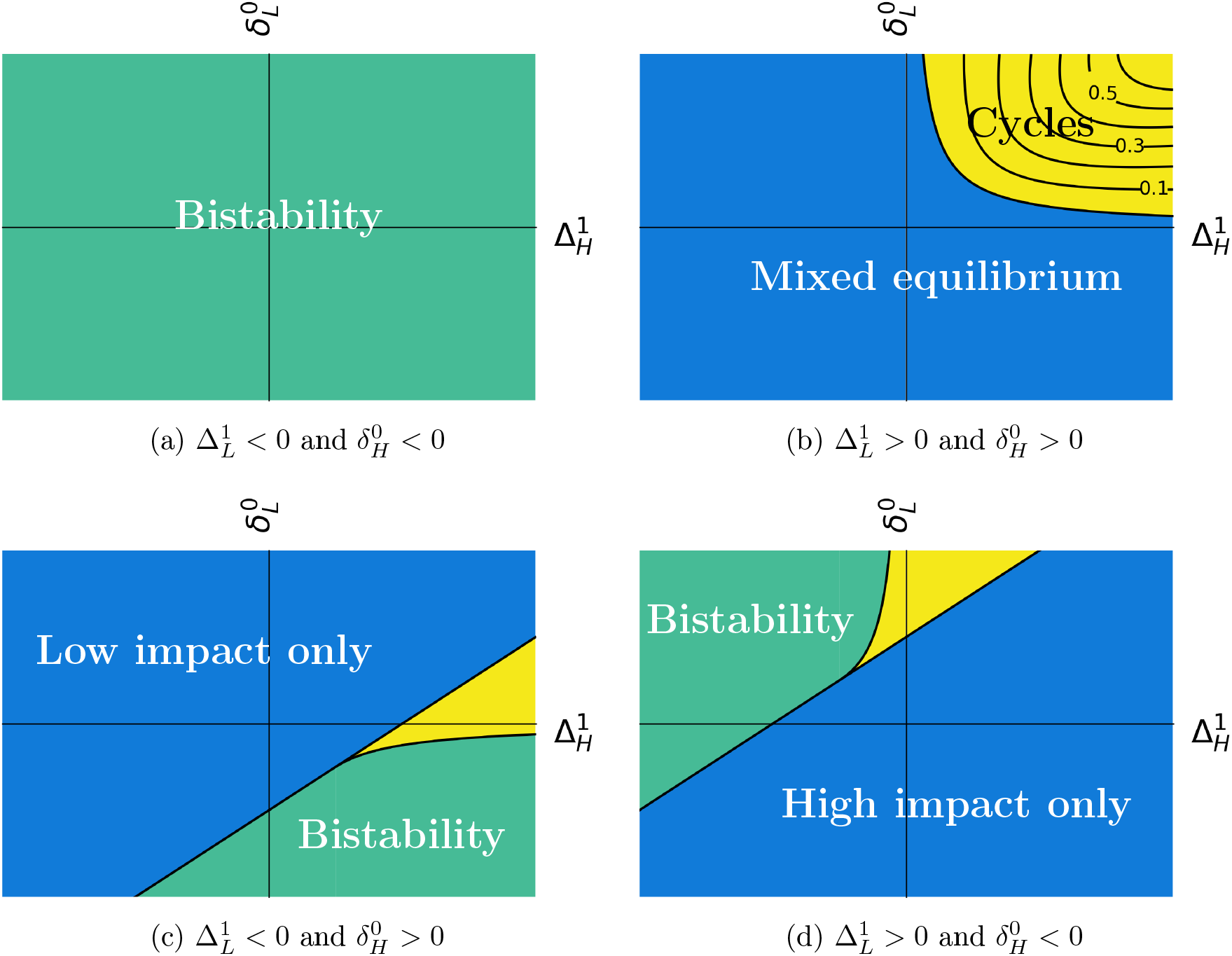
Dynamical outcomes of linear eco-evolutionary games. Each panel shows the dynamical outcomes for fixed values of incentives 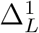 and 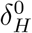. Yellow regions denote parameter regimes that can produce limit cycles, provided *ε < ε*_crit_, with level curves of ∊_crit_ shown as black lines. Blue regions represent regions with a single dynamical outcome. Green regions represent bistability. In panels (c) and (d) the yellow regions can exhibit bistability, dominance by one strategy, or cycles that occur in a bistable regime; the value of ∊ determines which of these outcomes occur. Supplementary Information section S1.2 provides analytical expressions for the boundaries between these dynamical regimes, in terms of the incentive parameters 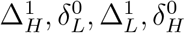.

When there are positive incentives for individuals to lead both the gold rush and the environmental movement (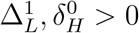; Figure 2b), then neither edge equilibrium is stable. In this regime, because individuals are always incentivized to lead change, environmental quality can possibly cycle over time. However, positive incentives to lead change are not sufficient to induce cycles. Cycles in this regime require that the incentives to be a follower of change are also positive and stronger, in aggregate, than the incentives to lead change. Furthermore, cycles also require that the environmental feedback is sufficiently slow compared to strategy evolution (Figure 3). In sum, when there are positive incentives to lead both movements, a stable limit cycle will occur when 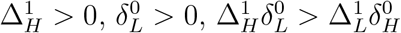, and *ε < ε*_crit_. We find no evidence of cycles outside this region (See SI S3).

**Figure 3:**
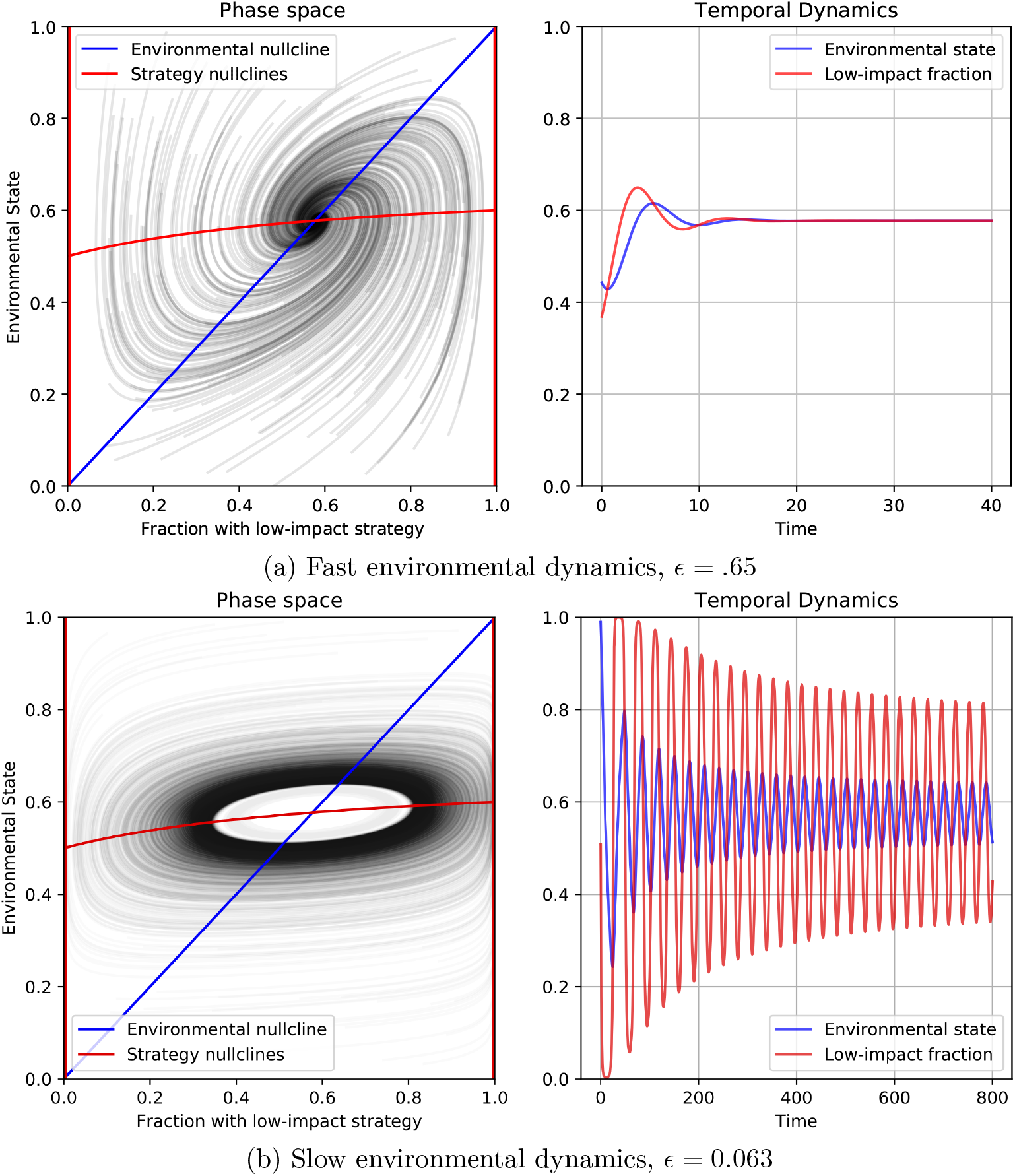
Phase planes and temporal dynamics in eco-evolutionary games with a decaying resource. Parameters are chosen to fall in the yellow region of figure 2(b). Only the speed of environmental dynamics relative to strategy dynamics, ∊, varies between the panels. Convergence either to an interior equilibrium or to a limit cycle is determined by ∊. 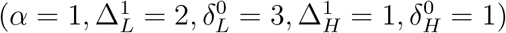

When there are positive incentives to lead the environmental movement but not to lead the gold rush (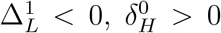; Figure 2 c), then a population composed of low-impact strategists will always be stable, whereas the high-impact strategic state will always be unstable. In this region we find parameter regimes that lead to a single monomorphic equilibrium, bistability, or stable limit cycles. In the blue-shaded region of figure 2 c there are no interior equilibria, so the system will settle on low-impact strategists alone. In the yellow and green regions, however, there are two interior equilibria. One of these is always a saddle while the other one can be stable or unstable. In the green region, with a stable interior equilibrium the system is bistable: it will approach either a population monomorphic for low-impact strategists, or a stable mix of both strategies. In the yellow region, the stable interior equilibrium becomes unstable under slow environmental feedbacks, leading to a limit cycle or a monomorphic population depending upon the rate of environmental feedback (Figure 4).

**Figure 4:**
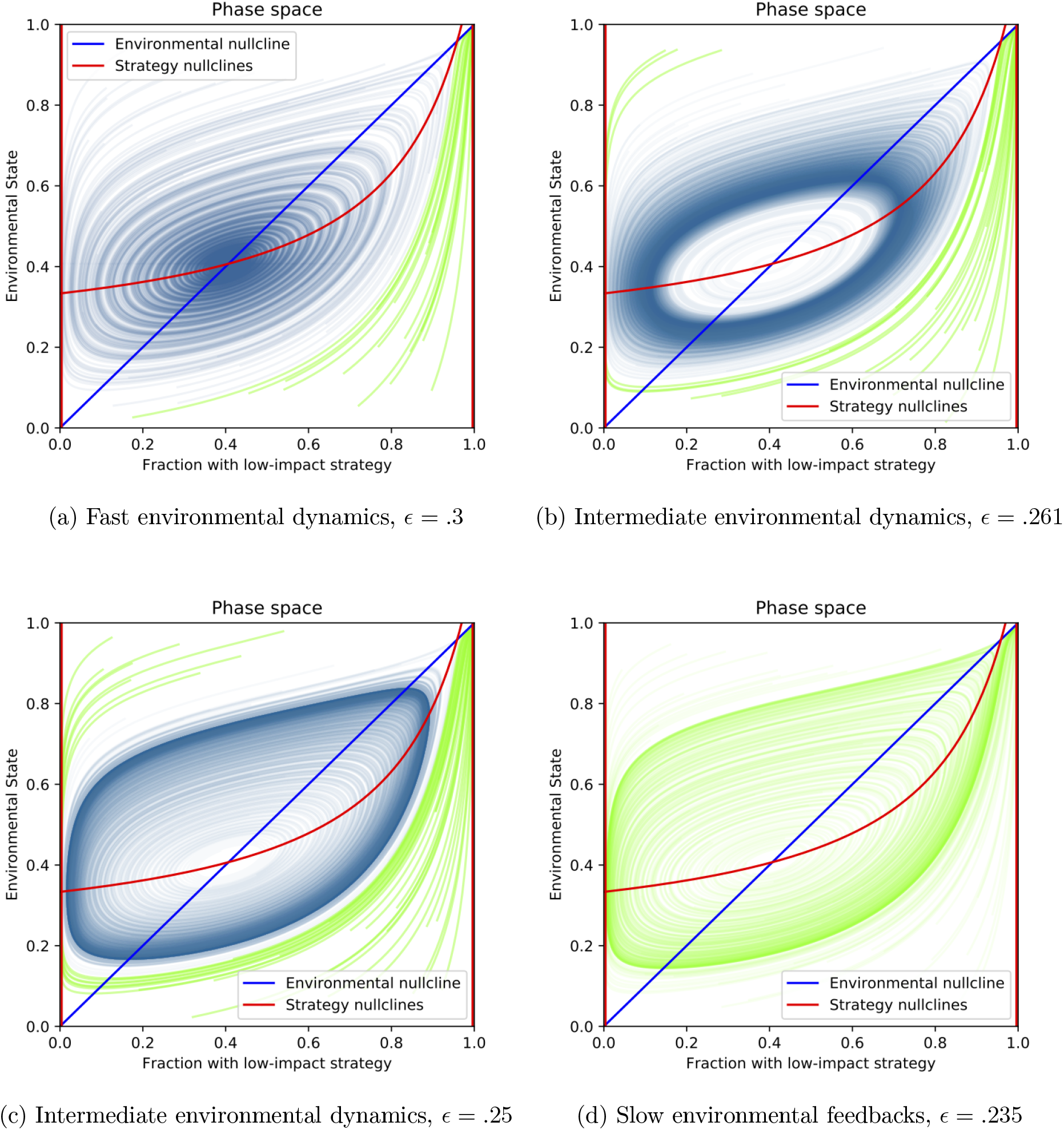
Simulations of eco-evolutionary game dynamics from the regime with possible cycles in figure 2(c). In each panel, the color of the solution curves illustrate the basins of attraction of the system. Green curves approach the state dominated by the low impact strategy and blue curves represent regions that approach the interior equilibrium or a stable limit cycle. Panel (a) shows the dynamics with fast environmental feedbacks, and thus the interior equilibrium is stable, and has a large basin of attraction. Panels (b) and (c) illustrate cases with environmental dynamics of intermediate speed which exhibit bistability between the low-impact dominated state and limit cycles in the interior of the phase space. Panel (d) displays dynamics under slow environmental feedbacks. When feedback speed crosses a critical threshold, limit cycles are no longer possible, and the entire phase space approaches the corner equilibrium. 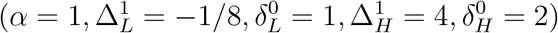

When there are positive incentives to lead the gold rush but not the environmental movement (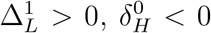; Figure 2 d), the high-impact state will always be stable and the low-impact state unstable. Here we find that, analogous to the regime in figure 2 c, bistability, limit cycles, and dominance by one strategy can all occur, depending on the relative incentives to follow change and the speed of environmental feedbacks.

The analysis above, catalogued by the four possible incentive regimes, gives a description of the possible outcomes for a linear two-strategy eco-evolutionary system (for a more detailed analysis see Supplementary Information section S1.2). The most striking result is that five easily interpretable parameters determine the qualitative properties of the system: four parameters that describe the incentives at the corners of state space, and one parameter describing the relative speed of environmental feedbacks. These parameters give immediate insight into when bistability, cyclic dynamics, mixed equilibria, or dominance by a single strategy can arise in eco-evolutionary games.

The speed of environmental feedbacks plays an important role in determining both the stability characteristics of equilibria and the basins of attraction of equilibria. Figure 4 shows the approximate basins of attraction for two equilibria under environmental dynamics of different speeds. These figures correspond to the region of potential cycles in panel (c) of figure (2).

Our analysis in this section has assumed that payoffs are linear in the state of the environment and the frequencies of the strategies in the population. These assumptions allowed us to characterize all possible outcomes in terms of a few parameters. While these simplifying assumptions may seem to limit the range of applicability, in the remainder of the paper we highlight scientifically and societally relevant cases that fall within this model, as well as examples that extend beyond the linear framework.

## 3 Case studies

A large collection of prior studies on environment-strategy feedbacks, across a range of disciplines, can be be understood as linear eco-evolutionary games. The dynamical properties of these models are all predicted by our analysis of these systems. In this section, we briefly review these models to highlight the broad applicability of our framework and to showcase the diversity of dynamical phenomena that occur in eco-evolutionary games.

### 3.1 Co-evolution of the environment and decision-making

Rand et al. (2017) developed a psychological model of decision-making where individuals can either make automatic “hardwired” decisions, or can make controlled decisions that are flexible and can shape a beneficial state of the environment. Although the motivation of their model is far from ecology, their formulation is a special case of the decaying resource model. Rand et al. found that under certain circumstances these feedbacks can lead to cyclical dynamics: automatic and controlled agents cycle in abundance as the environment fluctuates in its favorability towards these the two cognitive styles. Rand et al. (2017) motivate their study by noting that controlled decision making is likely to be costly but will allow individuals to choose optimal behavior. Further, they assume that when a population is dominated by controlled agents, then the fitness difference between optimal and suboptimal decisions will decrease because institutions or public goods created by controlled agents will stabilize the environment. This then favors automated agents who choose an option quickly without paying the cost of controlled processing. Rand et al. (2017) introduce a parameter that makes the cost of control frequency-dependent, so that when control agents are rare, it is more costly for them to ensure a stable environment.

The minimal model by Rand et al. is a special case of our decaying resource framework, as we prove in supplementary information section S4. By mapping their model onto our framework, the dynamical properties are immediately understood in terms of the incentive parameters of our analysis. In particular, we find that the Rand et al. model falls within panel (b) of Figure 2 – i.e. there are positive incentives to lead change. Embedding their model as special case of linear eco-evolutionary games allows us to show, for example, that cycles arise due to the frequency-dependent cost of being a controlled agent which assures that the model falls within the yellow region of Figure 2b. Further, we can compute the critical time lag that produces cycles between cognitive styles, coupled with environmental cycles (Supplementary Information section S4).

### 3.2 Grass-legume competition

Evo-evolutionary games provide a natural framework for studying competition between grasses and legumes. Many legumes form symbioses with nitrogen-fixing bacteria, allowing them to thrive in nitrogen-limited environments. Through time, however, some of the fixed nitrogen becomes available in the soil to nearby plants. In effect, the plant strategy of nitrogen fixation both frees plants from nitrogen limitation, and it generates an environmental feedback that increases the availability of nitrogen in the soil. Grasses, on the other hand, do not fix nitrogen. Competition between obligately nitrogen-fixing legumes and grasses can be modeled as a special case of our decaying resource framework. Here the environmental state corresponds to the degree that nitrogen is limiting, and the two strategies correspond to the species present in the system. We can thus write the relative abundance of grasses (the low nitrogen emission strategy) as *x* and legumes (the high nitrogen emissions strategy) as 1 *− x*.

We can determine the qualitative dynamics that will arise in grass-legume competition, based on the *δ* and ∆ parameters and the relative timescale of environmental versus strategy dynamics, ∊.

First, consider the grass-dominated state, with nitrogen limitation at its greatest (*n* = 1). In this state, we expect legumes to be able to invade because of the advantage of nitrogen fixation in a low-nitrogen environment. Thus we expect 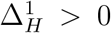. Similarly, we expect 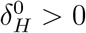 because in an environment where nitrogen is not limiting (*n* = 0) that is dominated by nitrogen fixing legumes, non-fixing grasses will be able to invade since they do not pay the cost of nitrogen fixation, but can reap the benefit of a nutrient-rich environment.

And so we know that the dynamics will fall somewhere in figure 2 b. Legumes are likely to have a competitive advantage in a low nitrogen environment regardless of their relative abundance, thus we expect 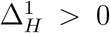. The same holds for grasses, given nitrogen is not limiting, so that 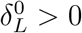. Finally, in species competition, it is typically more difficult for the first individual to successfully invade and establish than it is for an established species to spread and increase in abundance (Tilman, 2004). This implies that we expect 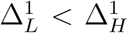 and 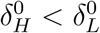.

Because 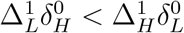 holds we expect grass-legume systems to be susceptible to cyclic dynamics. However, cyclic dynamics still require that the timescale of the feedback between the abundance of legumes and nitrogen availability is sufficiently slow. This too is reasonable in nature, because nitrogen is a valuable resource and a legume will tend to limit the rate at which fixed nitrogen leaks into the environment.

More broadly, feedback between plants and soil microbial communities, including through nitrogen fixation, can generate similar eco-evolutionary dynamics, with consequences for the maintenance of diversity (Bever et al., 1997; Mangan et al., 2010). A related feedback can occur between available soil nitrogen and nitrogen-fixation strategies of the rhizobium bacteria. Theoretical (Akçay and Simms, 2011) and empirical (Weese et al., 2015) findings indicate that high nitrogen availability favors rhizobia that fix less nitrogen, while low nitrogen favors strains that fix more. This suggests that cycling may also occur in coupled nitrogen-strain frequency dynamics.

### 3.3 Common-pool resource harvesting

Our final case study is a classic example of linked strategy and environmental dynamics: common-pool resource harvesting. There is extensive literature on common-pool harvesting, which forms the basis for bioeconomics (Clark, 1990). Eco-evolutionary game theory provides a natural framework to situate common-pool resource models – because strategic interactions depend upon, and conversely influence, the abundance of the common-pool resource.

It seems plausible that even the simplest form of common-pool resource harvesting will lead to cyclic dynamics: as the biomass of resource stock collapses and overshoots, harvesters respond by reducing effort, until the resource rebounds and high-effort strategies are again profitable. Nevertheless, our analysis shows that such cycles will never occur without additional complications. We formulate common-pool resource dynamics by assuming individuals can harvest with either high, *e*_*H*_, or low, *e*_*L*_, effort. We let the evolutionary process on strategy frequencies be governed by a profit function,

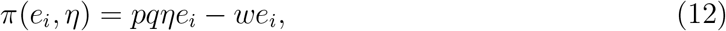

that maps the resource level and harvest effort into fitness. As in the general renewable resource model, we assume that *η* is governed by logistic growth, and that the harvest rate is proportional to *η* and effort (*e*_*L*_, *e*_*H*_). Since *π*(*e*_*i*_, *η*) is linear, this model is a special case of the renewable resource model we have exhaustively analyzed.

Transforming the resource level into a normalized environmental metric, we can construct a payoff matrix, Π(*n*) that maps the common-pool resource harvesting model onto our framework of eco-evolutionary games. The resulting parameter values satisfy 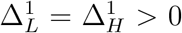 and 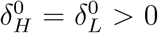 when there are positive profits at the environmental state resulting from pure low-impact strategists and negative profits at the environment resulting from pure high-impact strategists. Under these profit assumptions, the common-pool resource system falls at the boundary of the blue and yellow regions of Figure 2b – i.e. incentives to lead and follow change are all positive, but the there is no (positive) value of ∊ that produces cycles. And so the only possible outcome of this common-pool system is a stable mix of low- and high-impact strategists (see Supplementary Information section S5 for detailed analysis and for other possible scenarios). However, since the system falls on the boundary of a parameter region that permits cycles, small changes to the system may induce cyclic dynamics.

We considered two extensions, introducing market pricing (where *p* decreases as harvest quantity increases) or introducing harvesting efficiency gains (where *q* increases as a harvest strategy becomes more common). Both extensions fall outside the scope of the linear eco-evolutionary games analyzed in this paper. Market pricing induces non-linearity in the payoffs that harvesters receive, and frequency dependent harvest efficiency alters environmental dynamics outside of the renewing and decaying resource models considered above.

Under common-pool resource harvesting with market pricing, while the range of dynamical outcomes increases (See supplementary information figure S2), we do not find cyclic dynamics (see analysis in supplementary information section S6). This result occurs because market pricing effects all harvesters in the same way, and thus does not provide the extra incentive for being a follower of strategy change that can cause cycles.

Harvest efficiency may depend on strategy frequency if each strategy requires specialized skills and labor. As a strategy increases in frequency, increased opportunities for social learning may lead to increased proficiency and efficiency gains (Henrich, 2004; Smolla and Akcay, 2018). This effect alters both payoffs to individuals and the dynamics of the resource (see supplementary information section S7 for analysis). As a result of these intertwined consequences, we find instances of non-monotonicity – where increasing the growth rate of the resource, for example, can first destabilize and then stabilize an interior equilibrium (see supplementary information Figure S4). Despite this added complexity, the intuition developed from our general framework still applies. In particular, our analysis of common-pool resource harvesting as a linear eco-evolution game showed that increasing the values of either 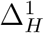 or 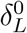 could cause cycles, by moving the system into the yellow region in Figure 2b. Frequency-dependent harvest efficiency plays a similar role to increasing 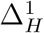 or 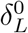, by making it more profitable to switch to a high-frequency strategy due to increased efficiency, and helps explain the cyclical dynamics that arise in this case (see supplementary information section S7).

## 4 Discussion

We have developed a framework for studying linked environmental and evolutionary game dynamics. Our analysis provides a systematic account of dynamical outcomes for an arbitrary game with linear payoff structures, for environments that either intrinsically grow or decay. This framework applies to any game-theoretic interaction where the strategies that individuals employ impact the environment through time, and the state of the environment influences the payoffs of the game. Such feedbacks are common in social-ecological systems, evolutionary ecology, and even psychology.

Despite the broad relevance of eco-evolutionary games, the role of the environment on strategic interactions has typically been considered only implicitly. And explicit account of environmental feedbacks reveals added complexity and nuance (Estrela et al., 2018). We have shown that many prior studies of strategic interactions with environmental feedbacks are special cases of our framework, with an explicit mapping to the parameters of our framework.

Perhaps the most striking result is that the rich dynamical outcomes that arise in ecoevolutionary games can be understood in terms of five intuitive parameters (Figure 2): the incentives to lead or follow strategy change under rich or poor environmental conditions, and the relative timescale of strategic versus environmental dynamics.

Our paper extends the model of Weitz et al. (2016) by incorporating ecologically inspired environmental feedbacks. The main difference between our work and Weitz et al. (2016) is that those authors consider an environment with no intrinsic tendency for growth or decay. In many systems of ecological and social relevance the environmental variables that affect payoffs have intrinsic dynamics, meaning that they regenerate (e.g., a population) or decay (e.g., pollution) when left by themselves. Our analysis shows that the choice of feedback structure has important consequences for system outcomes. Whereas Weitz et al. (2016) find heteroclinic cycles independent of the timescale separation between environmental and strategy dynamics, we find that the existence of limit cycles depends critically on the degree of timescale separation. And so the qualitative outcomes of eco-evolutionary games depend fundamentally on whether or not we account for the realistic intrinsic dynamics of an environment.

We have also seen that coupling strategies and the environment can induce persistent cycles even when neither the intrinsic environmental or intrinsic strategic dynamics exhibit cycles on their own. However, it is well known that many ecological systems can produce complex environmental dynamics even without feedback from individual actions (Turchin, 2003). Intrinsic complexity can result from multiple interacting environmental factors or a single environmental factor that is subject to age- or stage-structured population dynamics. In these cases, the complexity introduced via coupling with strategic interactions is likely to be even more rich, with the potential for chaotic dynamics.

Structured interactions, arising from environmental heterogeneity, network structure, and spatial structure will add further complexity. Spatial models of environmental feedbacks indicate that feedbacks can lead to correlations between strategies and their environment, with consequences for the stability of cooperation (Pepper and Smuts, 2002; Hauert et al., 2018). Spatially structured interactions, and more generally, network structured interactions among individuals can fundamentally alter predictions about the strategies that will be successful in a system, but it is not well known what effects such structured interactions will have on a system where strategies and the environment are coupled. Recent work suggests that it could lead to environmental heterogeneity and the simultaneous support of many strategies with differing environmental and strategic consequences (Lin and Weitz, 2019).

We focused most of our analysis on linear two-strategy eco-evolutionary games. And yet non-linearities in the payoff structure of strategic interactions have important effects. We have seen that introducing non-linear payoffs, through market pricing, in a common-pool resource harvesting model generates a broader class of qualitative outcomes, and the possibilities under a general non-linear payoff functions are likely even broader. Further, as the size of the strategy space increases, so does the dimensionality of the dynamical system. If new strategies emerge then the game itself can evolve (Worden and Levin, 2007), leading to novel interactions with the environment. These extensions make it clear that eco-evolutionary games can be seen as complex adaptive systems with emergent properties that are not easily predicted from the environmental dynamics or the evolutionary game structure alone.

In human societies the institutions structuring social interactions can be seen as part of the environment that co-evolve with strategic behaviors (Ostrom, 1990; Douglass, 1990; Bowles et al., 2003; Tilman et al., 2017). We can represent the institutional environment in our framework by constructing an institutional metric that modulates the game being played. Institutions may also interact with an explicit resource and affect the incentives individuals face in a given environment, or how strategies affect environmental dynamics. For example, the success of international environmental agreements to achieve environmental stewardship depends on how individuals and nations respond to their incentives under a changing environment. In the context of pollution control and climate change mitigation, action is an inter-temporal public good, with benefits of individual action accruing to a large population and in the future (Hauser et al., 2014). Policies of individual nations are likely to have feedbacks on the international institutional setting, the environment, and the choices that individuals make.

Environmental feedbacks in strategic interactions are the norm, not the exception. An explicit account of these feedbacks reveals commonalities among many societally relevant systems, ranging from the psychology of decision-making to species interactions and climate-change action, and alters predictions about expected outcomes in such systems. Incorporating strategy-environment feedbacks into evolutionary game theory is paramount in future studies.

## Supplementary Information

### S1 Renewable resource model

We analyze a two-strategy evolutionary-game-theoretic model that incorporates environmental feedbacks, governed by renewable resource dynamics. Suppose that there is a resource stock, *m*, that in the absence of consumption or harvest pressure grows logistically, and is diminished through harvesting or consumption that is associated with the strategies in a game. For a 2-strategy game, let *e*_*L*_ and *e*_*H*_ be the harvest effort of strategies L and H, respectively, where we assume, that *e*_*L*_ < *e*_*H*_. The dynamics of *m* are governed by

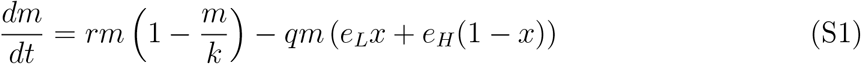

where *x* is the fraction of the population playing strategy L, *r* is the intrinsic rate of growth, *k* is the carrying capacity of *m*, and *q* is a parameter that maps resource degradation pressures (or harvesting efforts) (*e*_*L*_, *e*_*H*_) onto the rate of reduction in the resource. We assume that environmental impact rates are restricted so that *m* will be positive at equilibrium. This implies that *e*_*H*_ ∈ [0, *r/q*) and *e*_*L*_ ∈ [0, *e*_*H*_).

Let *n* ∈ [0, 1] be a normalized measure of the state of the environment that maps onto the payoff structure of the game. We relate *m* to *n* with the linear relationship

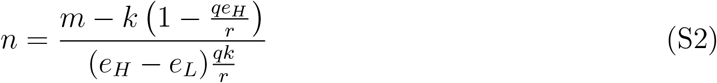

so that when the whole population chooses strategy L, corresponding to harvesting pressure *e*_*L*_, the equilibrium value of *m* maps to the environmental metric *n* = 1. Similarly, if the whole population adheres to strategy H, with *e*_*H*_ environmental pressure, *n →* 0.

Next, suppose that the state of the environment influences the payoffs of the game. We use a payoff matrix for a 2-strategy game with payoffs that are dependent on the normalized environmental measure, *n*, given by

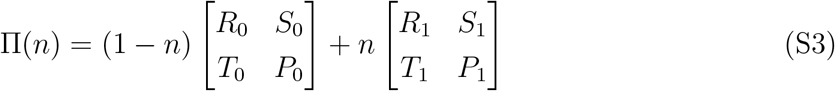

so that the matrix entries correspond to the payoffs of the game under conditions of a poor or rich environmental state. We can write the payoff for playing strategy L and strategy H as

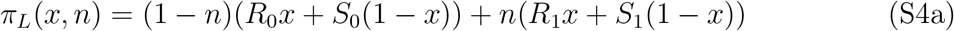

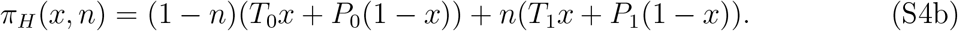

Following the replicator equation, the payoffs from the game determine the evolution of the fractions of the population that play each strategy. In all, we can write our system as

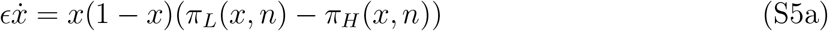

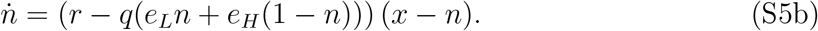

We let *τ* = *t/∊* and re-scale time so that our system can be written as

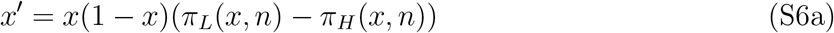

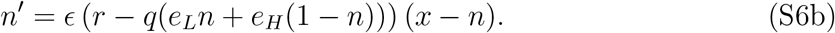

where 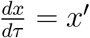.

#### S1.1 Renewable resource stability analysis

This model has up to four equilibria within the state space. We have 2 edge equilibria, at (*x^∗^, n^∗^*) ∈ {(0, 0), (1, 1)} and up to two interior equilibria that occur when *π*_*L*_(*x, n*) = *π*_*H*_ (*x, n*) and *n* = *x*. While there are up to two points that meet the criterion for a potential interior equilibrium, they need not fall inside the state space. Also, note that while *n′* = 0 at 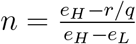 this too falls outside of our state space since we assume that *e*_*H*_ ∈(0*, r/q*) and *e*_*L*_ ∈ (0*, e_H_*).

At the edge equilibria, stability conditions are simple. At (*x^∗^, n^∗^*) = (0, 0), stability occurs if and only if *P*_0_ *> S*_0_. At At (*x^∗^, n^∗^*) = (1, 1), stability occurs if and only if *R*_1_ *> T*_1_.

Next, we analyze the conditions for stability of interior equilibria. The interior equilibrium, defined by *π_L_*(*x, n*) = *π_H_* (*x, n*) and *x* = *n* occurs when

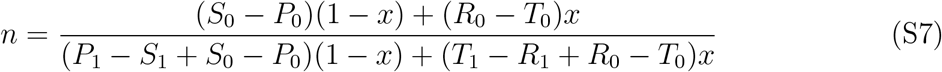

and *x* = *n*. We can substitute and solve for the equilibrium level of the environmental indicator and the equilibrium strategy fraction, which is

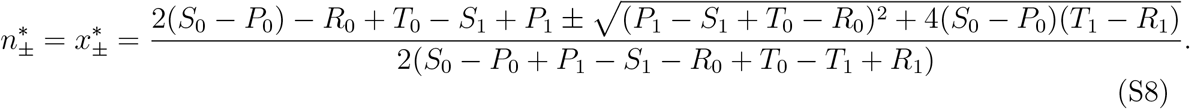

The analysis can be streamlined by defining four values as

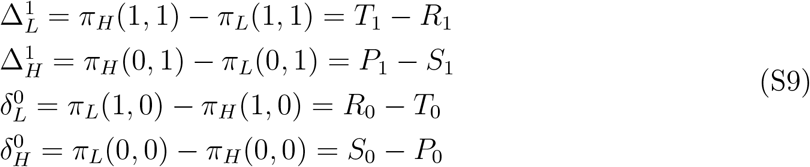

where the ∆’s correspond to the incentive to switch to the high impact strategy and the *δ*’s correspond to the incentive to switch to the low impact strategy. The superscripts correspond to the environmental state and the subscripts denote the resident strategy in the population.

Using these *δ*’s and ∆’s allows us to write the interior equilibrium location as

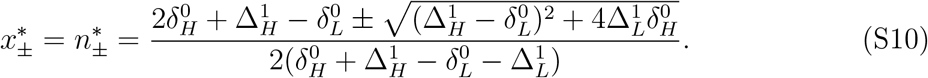

Local stability at such an equilibrium can be computed from the Jacobian matrix evaluated at an interior equilibrium, which is

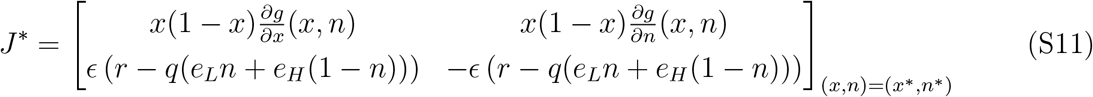

where *g*(*x, n*) = *π*_*L*_(*x, n*) − *π*_*H*_ (*x, n*). Stability requires Det(*J^∗^*) > 0 and Tr(*J^∗^*) < 0.

First, consider the determinant of the Jacobian matrix,

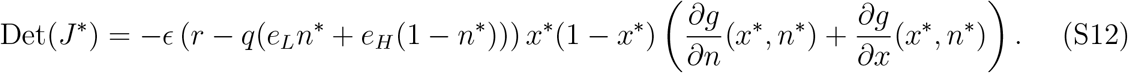

Due to the restrictions on the level of environmental impact, *e*_*H*_ ∈ (0, *r/q*) and *e*_*L*_ ∈ [0, *e*_*H*_), and the fact that both *x* and *n* are between zero and one, we can conclude that Det(*J^∗^*) *>* 0 if and only if 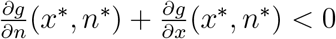. The terms of interest are

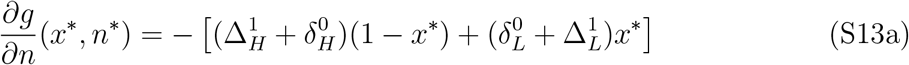

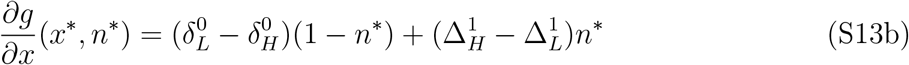

Recalling the at an interior equilibrium *x^∗^* = *n^∗^* we can simplify the expression for a positive determinant of the Jacobian matrix to

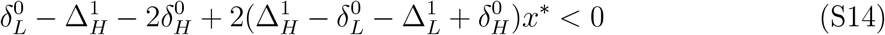

Evaluation at the equilibria yields the condition for a positive determinant, and is

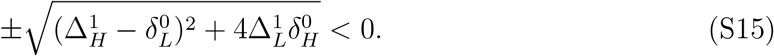

Where 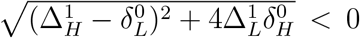 corresponds to 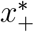 and 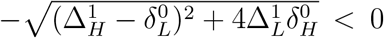 corresponds to 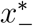.

The trace of the Jacobian matrix is

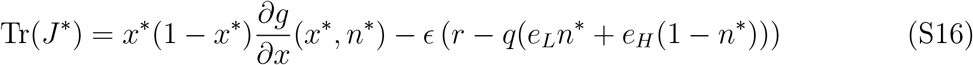

where *g*(*x, n*) = *π*_*L*_ (*x, n*) − *π*_*H*_ (*x, n*). Given Det(*J^∗^*) *>* 0, stability at an interior equilibrium will occur if and only if Tr(*J^∗^*) *<* 0. A sufficient condition for a negative trace is 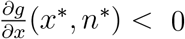, because this implies that both terms of the trace are negative. This occurs when either

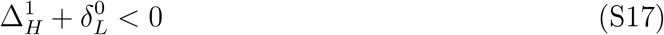

or

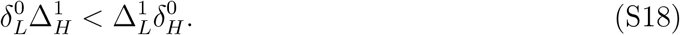

On the other hand if conditions S17 and S18 are violated, stability is still possible but not assured. In this scenario, stability depends on the relative speed of environmental feedbacks.

In this case, stability occurs at the interior equilibrium when

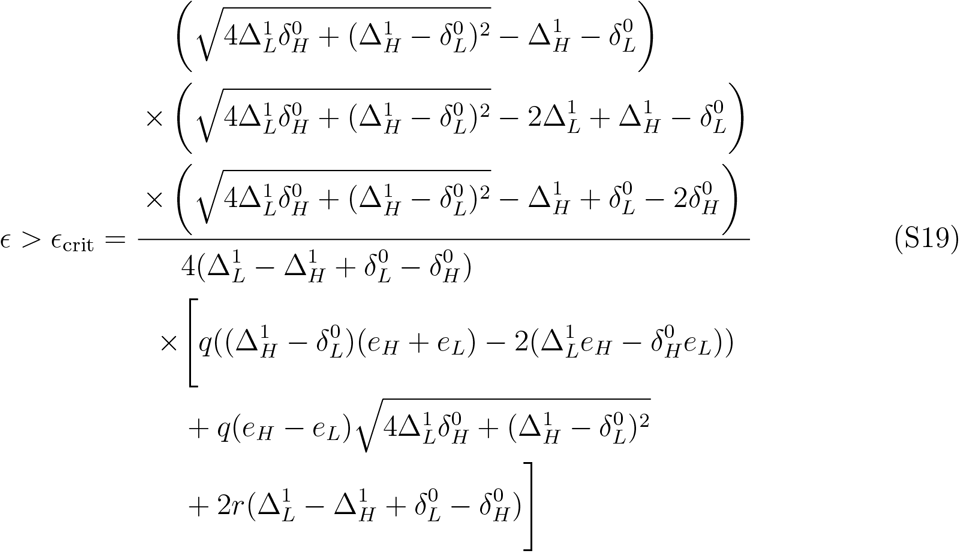

where ∊ is the speed of environmental feedbacks relative to strategy updating. As *r* increases, the region of the parameter space that leads to a stable interior equilibrium increases. This makes sense, both ∊ and *r* increase the speed of environmental dynamics, and thus changes in these parameters have similar effects on outcomes.

#### S1.2 Renewable resource system-level analysis

The 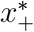 equilibrium is an element of the unit interval when the following conditions hold:

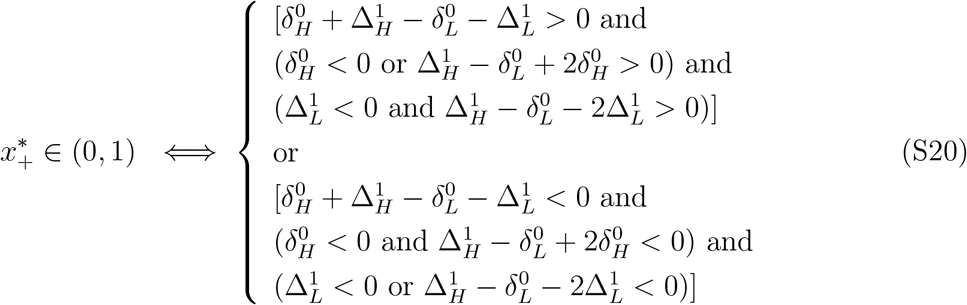

Similarly, the 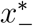 equilibrium is an element of the unit interval when the following conditions hold:

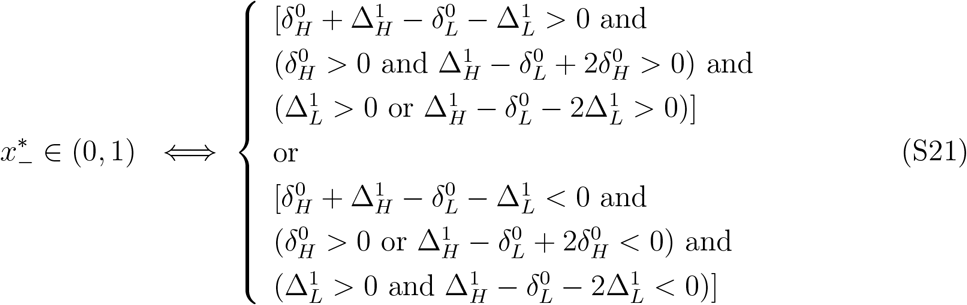

Ascertaining what these criteria mean for system-level properties of interest is simplified by considering four cases that span the values of 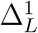 and 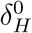.

##### Case 1: 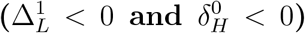

In this case, both edge equilibria are stable and only an interior saddle equilibrium, given by 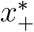 falls in the interior of the state space. Thus, this corresponds to bistability of strategy L and H where the resulting environmental and strategy state depends on the initial environmental and strategy conditions. This result is invariant to the details of the environmental feedback timescale or the payoff structure of the game (other than the restriction that 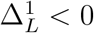 and 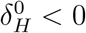).

##### Case 2: 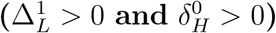

In this case both edge equilibria are unstable, and only the 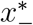 equilibrium is in the state space. The stability analysis indicates that 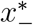 is guaranteed to be a locally stable equilibrium when either 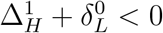 or 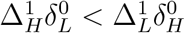. If both of these conditions are violated, then stability will governed by the condition on ∊. When the interior equilibrium is stable, we find no evidence limit cycles and expect all initial conditions to lead to the interior equilibrium. When it is unstable, limit cycles will result from all starting conditions. These are the cases that correspond to the ‘oscillating tragedy of the commons’ in Weitz et al. (2016).

##### Case 3: 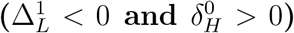

In this case, both interior equilibria can fall within the state space, given all the following hold:

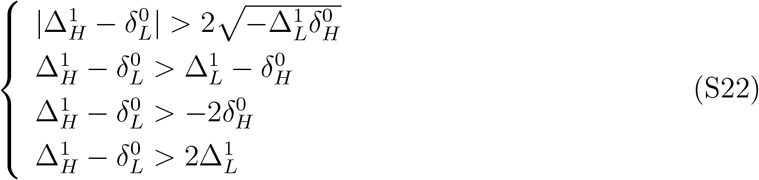

For these four conditions to hold, given 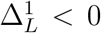 and 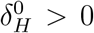, we need either 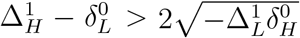 or 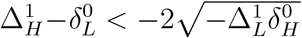. The latter cannot hold simultaneously with the three other conditions, since 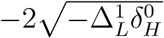 must be less that at least one of the other terms that set minimums on 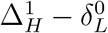. Further, the right-hand-side of the first inequality is positive, while the rest are negative. Thus all four of the conditions hold if and only if 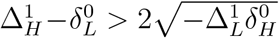. Also, since violating the first condition implies that interior equilibria cannot exist, there will either be two or zero interior equilibria (with a special case where the nullclines are tangent at one point). When there is no interior equilibrium, dynamics will tend toward the state where all individuals employ the strategy with low environmental impact. This state is stable, since 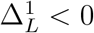. When there are two interior equilibria, one will be a saddle, and one will be stable or unstable depending on parameters and the degree of timescale separation. If this interior equilibrium is stable, then the resulting dynamics of the system will be similar to the bi-stable regime described above. If this interior equilibrium is unstable, the system may have limit cycles from some initial conditions and tend toward dominance of the low impact strategy from other initial conditions. It is also possible, for low values of ∊, that a limit cycle will not exist, and instead dynamics will tent toward the low impact state from all initial conditions.

##### Case 4: 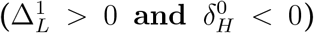

Lastly, following similar arguments as in case 3, we conclude that two interior equilibria will result when 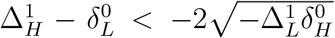, and no interior equilibria will occur if 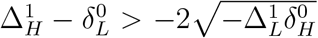. In this case, the edge equilibrium where the high impact strategy dominates will always be stable. If there are two interior equilibria, then as in case 3, depending on the value of ∊, either bi-stability, limit cycles embedded in a bi-stable regime, or dominance of the high impact strategy from all initial conditions will result.

### S2 Decaying resource model

While some resources are intrinsically self-renewing, many others are intrinsically decaying, and are maintained by production as a consequence of agents’ strategies. Here, we treat this case.

Let *m* be the concentration a resource that impacts the payoffs of players in a game, and is created as a byproduct playing the game. We assume that in the absence of production by players of the game, the concentration of *m* decays exponentially. We also assume that strategy L has a low emissions rate of the resource, *e*_*L*_, and strategy H has a higher emissions rate rate, *e*_*H*_. Given these assumptions we can model the dynamics of *m* as

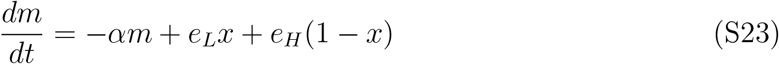

where *x* is the fraction of the population that employs strategy L, and (1 *− x*) is the fraction of strategy H players. While *m* can be any resource that meets the assumptions above, a clear example with societal relevance is pollution. Many actions generate pollution, and stocks of pollution impact many systems. However, there is a more broad class of problems that also fit within this framework, including the cognition-environment feedbacks studied by Rand et al. (2017).

As in the case of the regenerating resource studied in the previous section, we define a metric of the environmental state, *n*,

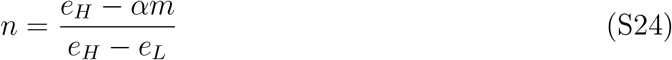

such that when the whole population has high emissions, *e*_*H*_, *n →* 0 and when the whole population has low emissions, *e*_*L*_, *n →* 1. Whereas *m* is a direct measure of the concentration of a resource stock, for example a pollutant, *n* is a normalized transformation of *m* that is used to write the payoff structure of the game being considered. Employing a change of variables, the dynamics of *n* can be modeled as

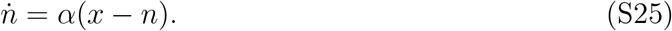

We use the same payoff matrix as before for the 2-strategy game with payoffs that are dependent on the normalized environmental measure, *n*, given by

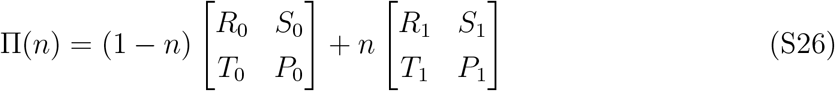

so that the matrix entries correspond to the payoffs of the game under conditions of a poor or rich environmental state. We can write the payoff for playing strategy L and strategy H as

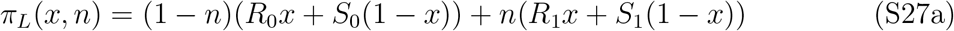

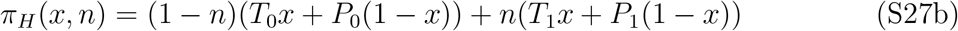

when *x* is the fraction of the population that plays strategy L and has low emissions. Following the replicator equation, the payoffs from the game determine the evolution of the proportion of the population that plays each strategy. In all, we can write our system as

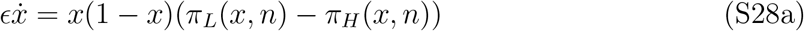

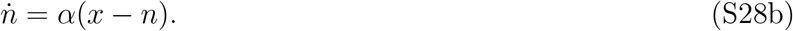

Let *τ* = *t/∊* to re-scale time so that our system can be written as

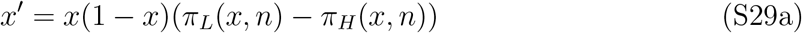

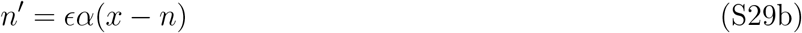

where 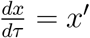.

#### S2.1 Decaying resource stability analysis

This model also has up to four equilibria within the state space. We have 2 edge equilibria, at (*x^∗^, n^∗^*) ∈ {(0, 0), (1, 1)} and up to two interior equilibria that occur when *π*_*L*_(*x, n*) = *π*_*H*_ (*x, n*) and *n* = *x*.

We analyze the Jacobian matrix to derive the conditions for stability of interior equilibria. The Jacobian matrix for this system at an interior equilibrium is

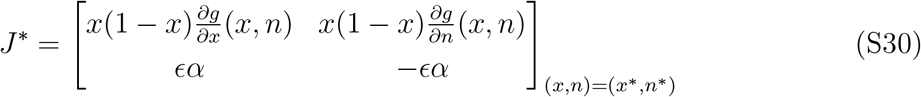

where, once again, *g*(*x, n*) = *π*_*L*_(*x, n*)−*π*_*H*_ (*x, n*). Also, the location of the interior equilibrium is the same as in the renewable resource model, with

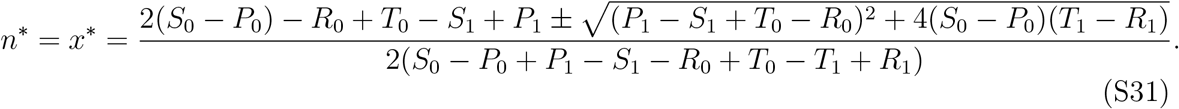

Again, we use

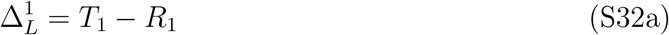

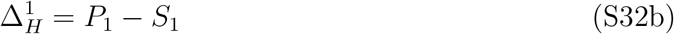

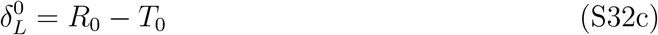

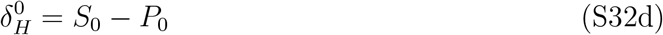

to simplify our expressions and take into account only the differences in the payoffs, without loss of generality. The simplified expression for the interior equilibrium is

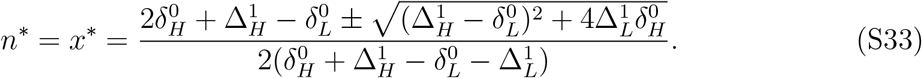

In relation to the analyses for the renewable resource model, the structure of the Jacobian matrix for the decaying resource model is very similar. This will make our analysis easier because much of it carries over directly from previous sections.

Stability at an interior equilibrium can be determined by trace and determinant of the Jacobian matrix. As before, the determinant is positive if and only if

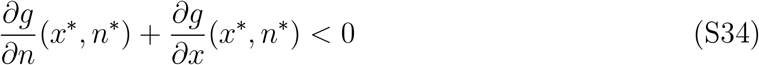

and a sufficient condition for a negative trace is

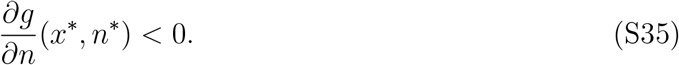

However, if condition S35 is violated but condition S34 holds, stability is still possible, and will depend on the speed of environmental feedbacks. In this case, the condition for stability is

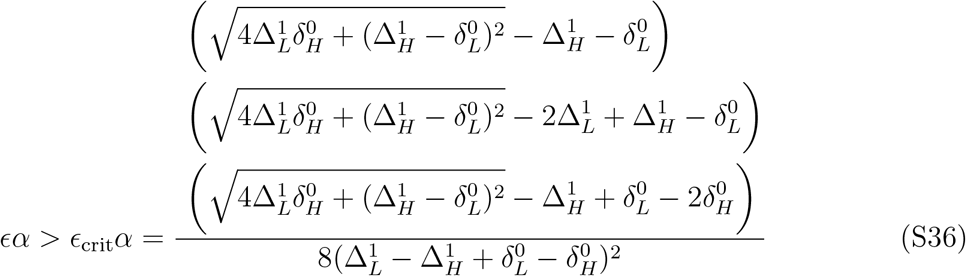

so that fast enough environmental dynamics can stabilize any interior equilibrium, even if condition S35 is not met. Lastly, *g*(*x, n*) is unchanged from the renewable resource model, implying that the forms of condition S35 and condition S34 also remain unchanged from the renewable resource model.

Now, we consider the edge equilibria. As before, at (*x^∗^, n^∗^*) = (0, 0) stability occurs if and only if 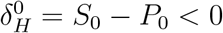. Again, (*x^∗^, n^∗^*) = (1, 1) is stable if and only if 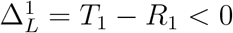. Thus if both 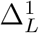 and 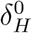 are positive, then both edge equilibria are unstable.

#### S2.2 Decaying resource system-level analysis

For the decaying resource model, the system-level analyses do not change from the renewable resource model, and thus the regions of the parameter space as divided in figure 2 do not change, and retain the same qualitative interpretations. This is because the position of the equilibria are unchanged and the sufficient conditions for stability of an interior equilibrium depend only on the *δ* parameters which share definitions in both systems. Thus, we can conclude that the only change to the system-level analysis from the renewable resource model is the value of ∊_crit_ that divides cycles from a stable interior equilibrium in the regions of the parameter space where cycles can occur.

### S3 The existence of cycles

We alluded the the existence of limit cycles in some regions of the parameter space, and suggested that limit cycles do not occur in other regions of the paratmeter space. A challenge to making such claims arises because local stability analyses are not sufficient to describe global dynamics. In this section we detial the analysis on the existence, (or non-existence) of limit cycles in our framework. We break this analysis down into cases, following our local stability analysis.

#### Case 1: 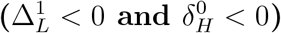

We showed that in this case the only interior equilibrium point is a saddle. In two dimensional systems, a limit cycle must contain at least one equilibrium point. However, in this region, the interior equilibrium is a saddle point, which cannot be the only equilibrium point within a limit cycle in a two dimensional system. Therefore, we can conclude that this region of parameter space cannot contain limit cycles.

#### Case 2: 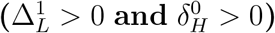

In this region only one equilibirum is present in the interior of the state space, and it is either stable or unstable, never a saddle point. Our local stability analysis indicated that when 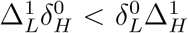 then decreasing ∊ below ∊_crit_ would destabilize the interior equilibirum, leading to super-critical Hopf bifurcation. Since in this setting there are no stable equilibria (in the interior or boundary of the state space) persistent oscillations result. Simulations indicate that these oscillations are stable limit cycles. While we can prove the existence of these oscillations for part a region of the parameter space, it is our goal to preclude them elsewhere.

Consider the region of the parameter space where 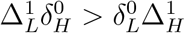. In this region, the slope of the strategy nulllcline, is negative. Thus, we can see graphically that for a fixed value of *n* dynamics converge to the strategy nullcline. This implies that when strategy dynamics are fast relative to environmental dynamics (small ∊), the interior equilibrium is globally stable since dynamics quickly approach the strategy nullcline, then slowly converge to the interior equilibrium. Conversely, if environmental dynamics are fast, dynamics quickly converge to the environmental nullcline then slowly converge to the interior equilibrium (whether 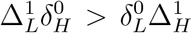 holds or not). Therefore, we can conclude that if limit cycles exist in the region of the parameter space where 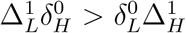, they must only occur for intermediate values of ∊.

Limit cycles can be ruled out by the Dulac-Bendixson theorem. If we can find a suitable function *φ*(*x, n*) such that

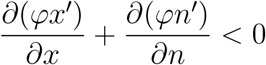

for all values of *x* and *n* in our state space, then we can rule out limit cycles for that region of the parameter space. Consider the Dulac function 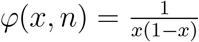 for the decaying resource framework. Then we seek to show that

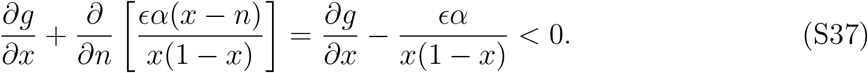

We are interested in the region of the state space for which limit cycles can be ruled out for any value of ∊. Notice that the term 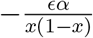 is always negative, but becomes close to zero for small ∊ and *x* near 1*/*2. Therefore, limit cycles can be ruled out when 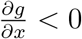 for all *x*, *n* in the state space. Thus we can rule out cycles for any ∊ when 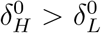 and 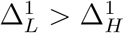. We can also rule out cycles when

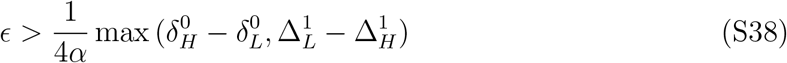

holds. This proof of the non-existence of limit cycles shows that limit cycles cannot occur for fast environmental feedbacks. Further, a graphical argument shows that limit cycles cannot occur for slow environmental feedbacks either (given 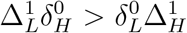, the fast sub-system is attracting and slow dynamics along the strategy nullcline lead to the equilibirum). Further, when 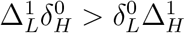 holds, then we know that the interior equilibrium will be locally stable for all ∊, which implies that if limit cycles were to arise in this region it would have to be a pair of unstable and stable limit cycles. In extensive simulations we find no evidence of such dynamics. For the renewable resource framework, the same analysis holds, given *e*_*L*_ > 2*e*_*H*_ − *r/q*.

#### Case 3: 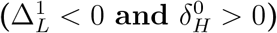

In this case there are either zero or two interior equilibria. In the cases where no interior equilibria are present, limit-cycles cannot occur, since a limit cycle must contain an equilibrium. In the region of the parameter space where there are two interior equilibria, one is always a saddle and the other is stable or unstable. The same analysis as above, in consultation with figure 2 shows that limit cycles cannot occur in this regime for 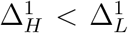. Further, any limit cycle that does exist will not contain the saddle equilibrium. While this is a proof of the non-existence of limit cycles for only a subset of the green region in figure 2, we again find no evidence of limit cycles in the green region of the parameter space.

#### Case 4: 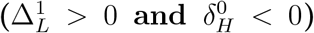

Lastly, following similar arguments as in case 3, we conclude that limit cycles cannot occur when 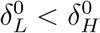.

### S4 Cognitive strategy dynamics as an eco-evolutionary game with decaying resource

Rand et al. (2017) present a model of automatic versus controlled processing with a cognition-environment feedback. In their model, the presence of controlled agents leads to an environment that favors automatic processing, and controlled processing is costly. They show that when the cost of control increases when there are fewer control agents, then cycles can result in what they term an “evolutionary pendulum”. The minimal model presented in their paper is a special case of our decaying resource framework of game-environment feedbacks. In this section, we make the connection explicit.

First, we introduce the model from Rand et al. (2017). Let *x* be the fraction of controlled processing agents in the population. Then the fitness of each agent is

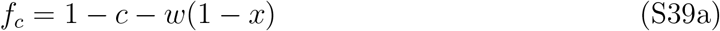

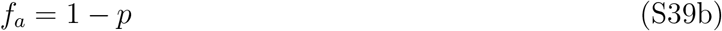

where *c* is the fixed cost of controlled processing, *w* is the extra cost of being a controlled processing agent when rare, and *p* is the environmentally dependent cost of automatic processing. This cost is governed by a “cognition-environment” feedback given by

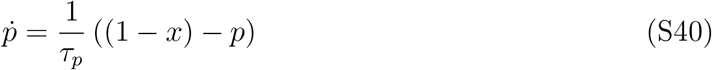

where *τ*_*p*_ governs the relative speed of this feedback. The dynamics of the frequency of the agents is governed by the replicator equation:

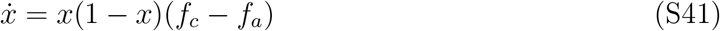

This leads a complete system of equations given by

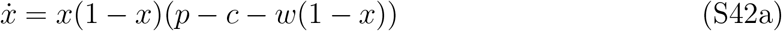

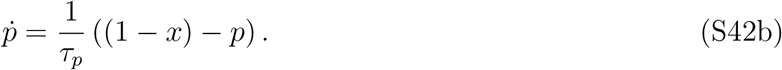

Recall that the general decaying-resource model presented herein is given by

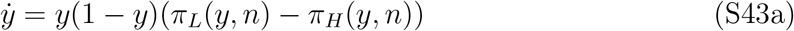

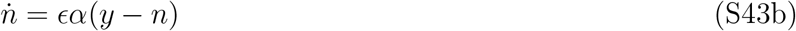

where *y* is the fraction of the population with strategy L, *n* is a measure of the resource stock and *π*_*L*_(*y, n*) and *π*_*H*_ (*y, n*) are the environmentally dependent payoffs from the game matrix

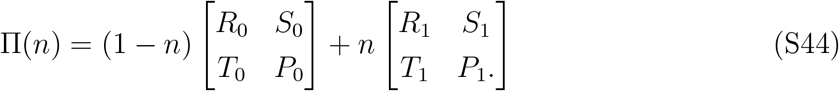

If we transform the equations from Rand et al. (2017) by

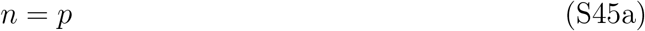

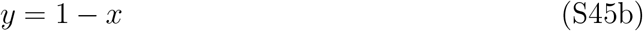

we can write the dynamics of the new system as

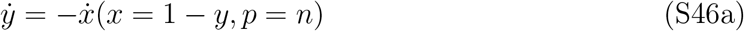

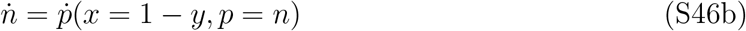

which simplifies to

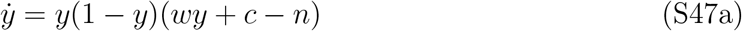

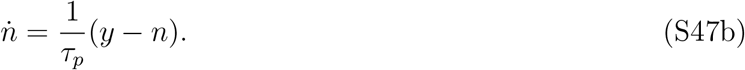

Thus, if we let *∊α* = 1/*τ*_*p*_, *R*_0_ = 1, *S*_0_ = 1, *T*_0_ = 1 *− c − w*, *P*_0_ = 1 *− c*, *R*_1_ = 0, *S*_1_ = 0, *T*_1_ = 1*−c−w*, and *P*_1_ = 1*−c* we can see that our general model from system of equations S43 becomes the same as a linear transformation of the Rand et al. (2017) minimal model. Thus the Rand et al. (2017) model is a special case of the decaying-resource model presented and analyzed in this paper. Given this, we could typically compute the *δ*’s and from this know the location, stability and dynamics near the equilibria of the system. However, this model is a ‘special’ special case, thus more analysis is needed to go from the general model to the results of Rand et al. (2017).

The *δ* values that govern the analysis are

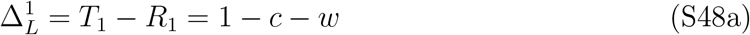

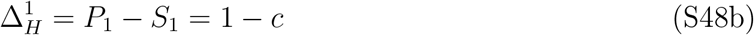

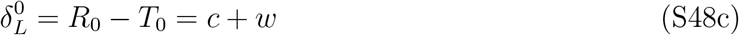

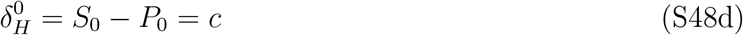

these values for the *δ* and ∆ parameters fall within the special case where

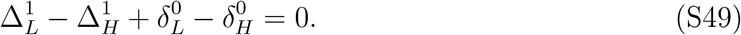

This is a special case that we have yet to analyze for the general model, because it accounts for a small region of the parameter space. We proceed to analyze the general model given we are in this special case. We are interested in the stability of the interior equilibrium because this is the equilibrium point around which the cyclic dynamics occur in the Rand et al. (2017) model.

#### S4.1 Analysis

Assuming that 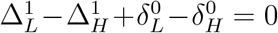 we analyze the general model with a decaying resource, given by

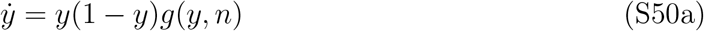

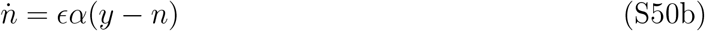

where *y* is the fraction of the population with strategy L, *n* is a measure of the resource stock and

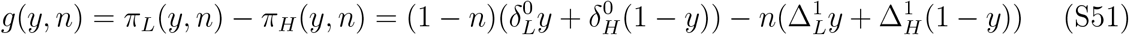

is the gradient of selection. The interior equilibrium occurs where *g*(*y, n*) = 0 and *y* = *n*. The first condition holds when

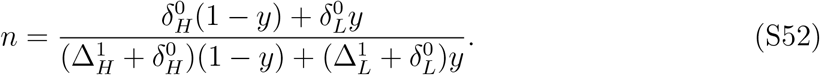

In this special case, the denominator is equal to 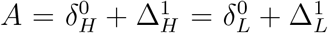. This simplifies our analysis and we can write the location of the interior equilibrium as

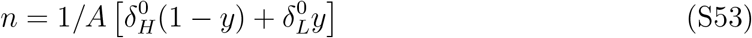

which, implies that

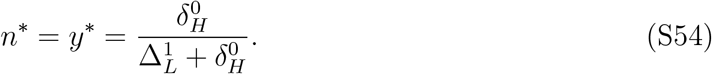

Stability at this interior equilibrium can be derived from the Jacobian matrix evaluated at this equilibrium,

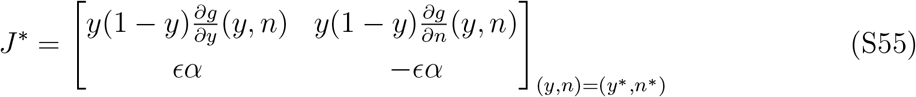

where

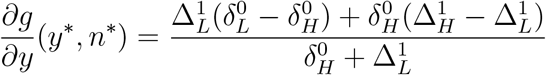

and

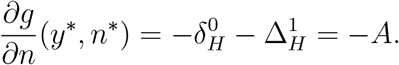

An unstable interior equilibrium will occur when Det(*J^∗^*) *>* 0 and Tr(*J^∗^*) *>* 0, and this will lead to cycles. Det(*J^∗^*) *>* 0 if and only if 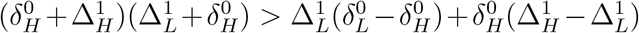, which in this special case is equal to

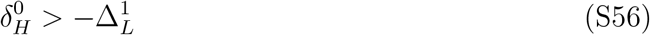

by the definition of *A*. Tr(*J^∗^*) *>* 0 if and only if 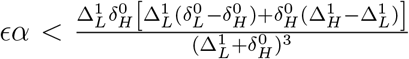 which in this special case is equal to

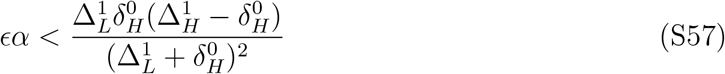

Now That the analysis has been done for the general case under the assumption that 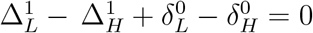, we can apply it to the model of Rand et al. (2017).

#### S4.2 Application to Rand et al. model

In the transformed version of the Rand et al. (2017) minimal model we had

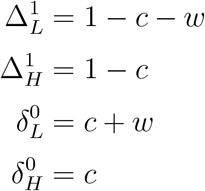

which can be applied to the conditions for an unstable interior equilibrium in equations S57 and S56. An unstable interior equilibrium occurs when

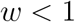

and

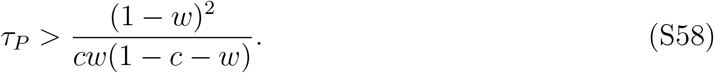

The first condition is satisfied by the assumption needed for the existence of an interior equilibrium, *w* + *c <* 1 and the assumption that *c* ≥ 0 and *w* ≥ 0. Notice that when *w* = 0, there does not exist a *τ*_*p*_ for which the interior equilibrium is unstable. This can be seen directly from a necessary condition for cycles, violating equation S18 which requires *w* > 0 (this is from the renewable resource case, but an identical necessary condition exists in the decaying resource case).

## S5 Common-pool resource harvesting

Here, we consider a classic model of common-pool resource harvesting. We assume that individuals either harvest with high, *e*_*H*_, or low, *e*_*L*_, effort. We let the evolutionary process be governed by a profit function, *π*(*e*_*i*_, *η*), that maps the resource level and harvest effort into fitness. As before, we assume that the resource, *η* is governed by logistic growth, and the harvest rate is proportional to *η* and effort. We can write our system as

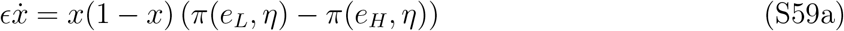

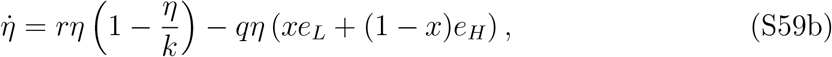

where *x* is the fraction of the population that harvests with low effort, *k* is the carrying capacity of the resource, *q* is the efficiency with which effort is transformed into harvest, and ∊ controls the relative timescales of strategy and resource dynamics. For small ∊, strategy dynamics are fast relative to resource dynamics.

Now, we turn our attention the the profit function, *π*(*e*_*i*_, *η*). In this simplest case, revenue will depend on the harvest and the price at which the harvested resource can be sold, and costs will increase linearly in effort. Under these assumptions, we can write the profit of an individual as

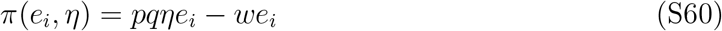

where *p* is the fixed price of the resource and *w* is the marginal cost of effort.

This gives us a complete description of our system, however, it is not immediately obvious that this model is a special case of the renewable resource case studied in detail above. Employing this profit function, we can simply express our system of equations as

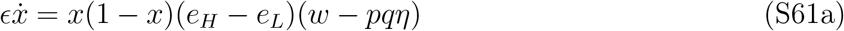

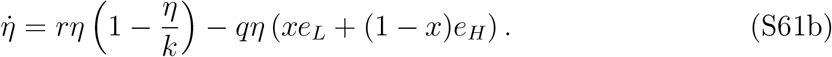

Throughout this analysis, We assume that *e*_*L*_ < *e*_*H*_. Now, in order to make an exact mapping between this model and the general renewable resource model analyzed in section 2.1 we linearly transform the resource level, *η*, into an environmental metric, *n* that is bounded between 0 and 1. Within this transformed space, we can construct a payoff matrix, Π(*n*) that maps onto our common-pool resource harvesting case.

First, we define our environmental metric as

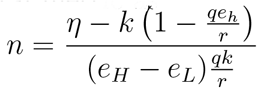

and write the state of our resulting system in terms of *x* and *n*. We have

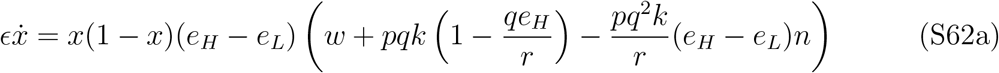

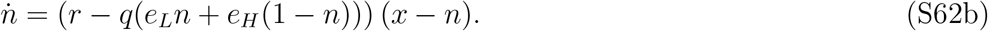

From this we can reconstruct a payoff matrix Π(*n*) that leads to this system equations. The entries of Π(*n*) are

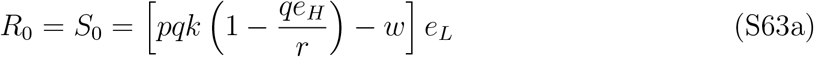

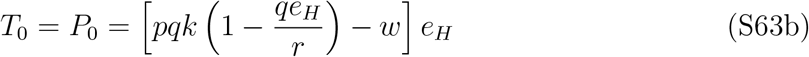

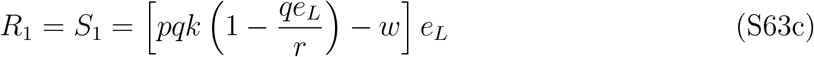

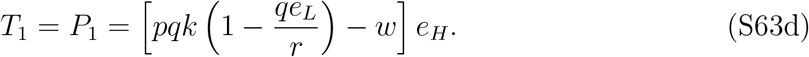

Thus, we can write the values of *δ* as

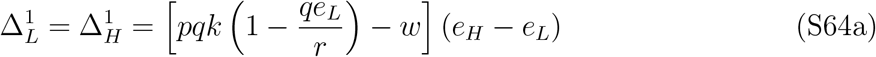

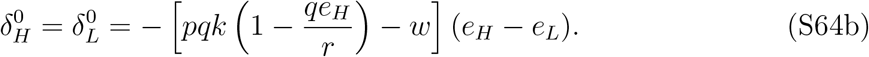

Depending on the parameter values chosen we have three possible outcomes. First, when positive profits occur at the low effort dominated environmental equilibrium but negative profits result at the high impact dominated equilibrium then we have 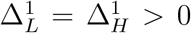 and 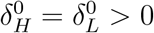. This implies that the system always falls at the boundary of the region where cycles are possible in Figure 2b. This implies that while cycles cannot occur in this system, small changes to the payoff structure could lead to cycles. In particular, holding all else equal, increasing 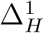 or 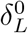 would permit cycles.

Second, if *e*_*H*_ is low enough that 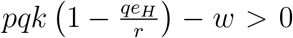, then 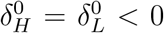 and 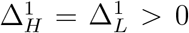. Thus, by referencing figure 2d, we conclude that the only stable outcome is the high-impact equilibrium. Third, if both 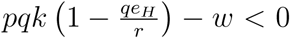 and 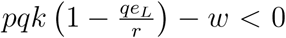, then we have 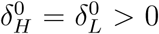 and 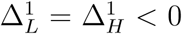. As illustrated in figure 2c, this implies that the low impact dominated state will be the equilibrium outcome. We assumed ex-ante that *e_L_ < e_H_* and thus these are the only possible outcomes within this system.

## S6 Market pricing CPR model

While the simplest common-pool resource harvesting model cannot produce cyclic dynamics, one possible driver could be market pricing. Whereas in the previous section the price received for each unit of harvested resource was constant, here we let the price received be a function of the total harvest at the present time, to mimic market pricing. A fixed price makes sense in the case where the system in question represents a small part of a large or global market because then local supply can have only a small impact on the global market price. Here, we allow harvest quantity to effect the price. This is more applicable to cases where the markets for the resource are local or the suppliers in consideration account for a significant portion of total supply. As a first approximation, we assume that the market price is linearly decreasing in the quantity supplied.

This alters that profit function, resulting in

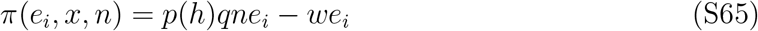

where *h* = *nq*(*xe*_*L*_ + (1 − *x*)*e*_*H*_) and *p*(*h*) is the market price as a function of resource supply.

This model is not a special case of the model presented in section 2.1 since there are higher order terms in the dynamical equations introduced through the dependency of the price of the resource on the state of the system. We consider the linear price equation,

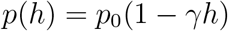

so that there is a price ceiling of *p*_0_ and a linear decrease with slope *γ* as supply increases. With this formulation, we can write the profit of low and high effort harvesters as

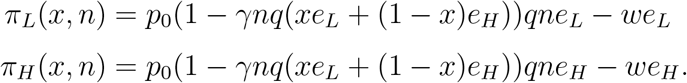

In total, the model can be written as

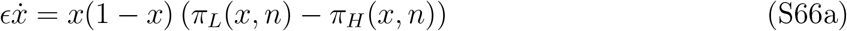

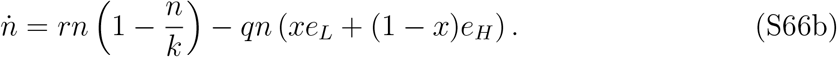

Bifurcation analysis allows us to show how equilibria (and their stability) change as a function of key parameters of interest. First, we examine the effect of the slope of the price function on the equilibrium fraction of low-effort harvesters. this slope, *γ*, is a measure of how quickly price declines as supply increases and is related to the price elasticity. The equilibrium state of the resource can also be displayed, however, in our model there is a linear relationship between the fraction of low-effort harvesters and the equilibrium state of the resource. Thus, we only present the bifurcation diagram in the fraction of low-effort harvesters, *x*. Figure S1a shows that under regime with a high maximum price, *p*_0_, increasing the rate at which harvest quantity depresses market price, *γ*, leads to hysteresis and multiple stable states. On the other hand, figure S1b shows that under a low maximum price, increasing *γ* leads to a smooth transition from one equilibrium to the other, without a region of multiple stable states.

### S6.1 Market pricing analysis

The interior strategy nullcline of the system changes from previous models because market pricing of harvest changes the set of points where zero profits result. Letting *π*(*e*_*L*_, *x, n*) − *π*(*e*_*H*_, *x, n*) = 0 and solving for *n* gives

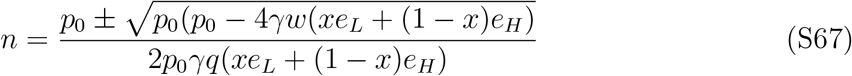

so that there are two branches of the relationship depending on whether the square root is added or subtracted from the relationship. This can be simplified to

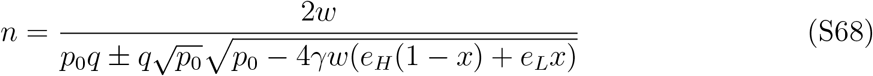

which, since *e*_*H*_ > *e*_*L*_, illustrates that as *x* increases, the second term in the denominator also increases. This implies that on the upper branch of the nullcline is increasing in *x* and the lower branch of the nullcline is decreasing in *x* for all *x* ∈ [0, 1], given that *p*_0_ *>* 4*γw*(*e*_*H*_ (1 − *x*) + *e*_*L*_*x*). This will simplify stability analyses at interior equilibria.

Now we characterize the stability of the equilibria of the system. First, we will consider the interior equilibria.

#### Interior equilibria

At interior equilibria, the environmental nullcline intersects with the strategy nullcline. The analysis will depend on the relative slopes of these nullclines, and whether the intersection occurs at the upper or lower branch of the strategy nullcline. Consider the Jacobian matrix

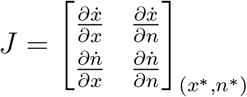

at the lower branch of the strategy nullcline. We know that the environment nullcline is increasing in *x* because *e*_*L*_ < *e*_*H*_, and that 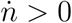 below the nullcline. Further, from the form of the strategy nullcline, we know that the lower branch is decreasing in *x* and that 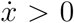 below the nullcline. From this we can determine the sign of each element of the Jacobian matrix. In this case we have

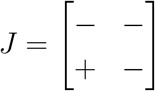

for the signs of the Jacobian, which implies stability.

Along the upper branch of the nullcline, the stability analysis is slightly more complex. Consider the path derivatives along the 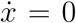 and 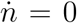 nullclines. By construction, the value of the path derivative is equal to zero, thus we can write

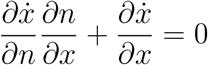

for the path derivative along the strategy nullcline where the slope of the nullcline is 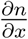. Similarly, we can write the value of the path derivative along the environment nullcline as

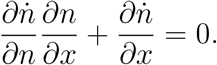

Let *S*_*x*_ and *S*_*n*_ denote the slopes of the strategy and environment nullclines at equilibirum, respectively. Using these identities, we can rewrite our Jacobian as

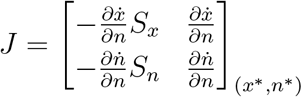

since *S*_*n*_ is positive and 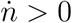 below the environment nullcline then 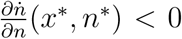. Also, since *S*_*x*_ is positive and 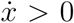 above the upper branch of the strategy nullcline, then we can conclude that 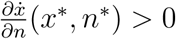. In total, this implies that Tr(*J*) *<* 0 and

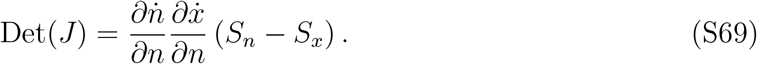

We can conclude that the interior equilibrium at an intersection of the upper branch of the strategy nullcline with the environment nullcline will be stable if and only if *S*_*x*_ > *S*_*n*_. If the slope of the environment nullcline is greater than the strategy nullcline, then the interior equilibrium at this point will be a saddle. Note that never does the stability of an interior equilibrium depend on the relative timescale of resource and strategy dynamics, ∊.

### Edge equilibria

Here, we analyze the stability of the equilibria at the edge of the phase space. These analyses are simplest graphically. Given that *e*_*L*_ < *e*_*H*_ and *e*_*L*_ is low enough that it will not drive the environment to zero, then both equilibria at *n* = 0 will be unstable. The equilibrium at *x* = 0 and *n* > 0 is stable if and only if it’s location falls above the lower branch of the strategy nullcline, but below the upper branch of the strategy nullcline. Conversely, the equilibrium at *x* = 0 and *n* > 0 is stable if and only if its location falls below the lower, or above the upper branch of the strategy nullcline.

Figure S2 shows a panel of eight qualitatively distinct phase diagrams of our model of market pricing, harvest and environmental dynamics. While there are many possible qualitative outcomes, none involve cyclic dynamics.

## S7 Frequency dependent harvesting efficiency CPR model

In this section we extend the common-pool resource model introduced in section 3.3 with the addition of a term to account for the dependency of harvest efficiency, *q*, on the frequency of a strategy. This model is not a special case of the general renewable resource model outlined in section (2.1) because the frequency dependent harvest efficiency terms alter the dynamics of the resource. The payoff function, however, is a special case of the payoff matricies considered in section (2.1). Suppose that low effort harvesters share information about the state of and location of the resource. This could lead to increasing harvest efficiency, *q*, as a function of the fraction of the population who harvest with low effort, *x*. Alternatively, each strategy may require specialized skills and labor. As a strategy increases in frequency, the supply of these skills increases, potentially leading to efficiency gains. In figure S3 dynamics are shown for the case that only low effort harvesters have efficiency that increases in their frequency. This can lead to limit cycles depending on the relative speed of the resource dynamics. We formalize this by considering a catchability function, *q*(*x*) that increases linearly in *x* for low effort harvesters, and decreases in *x* for high effort harvesters. This indicates that when a particular strategy is more common, the harvest of the resource by players with that strategy becomes more efficient. This could result form communication about the location of the resource, which would lead to an increase in harvest per unit effort for a given environmental state.

**Figure S1:**
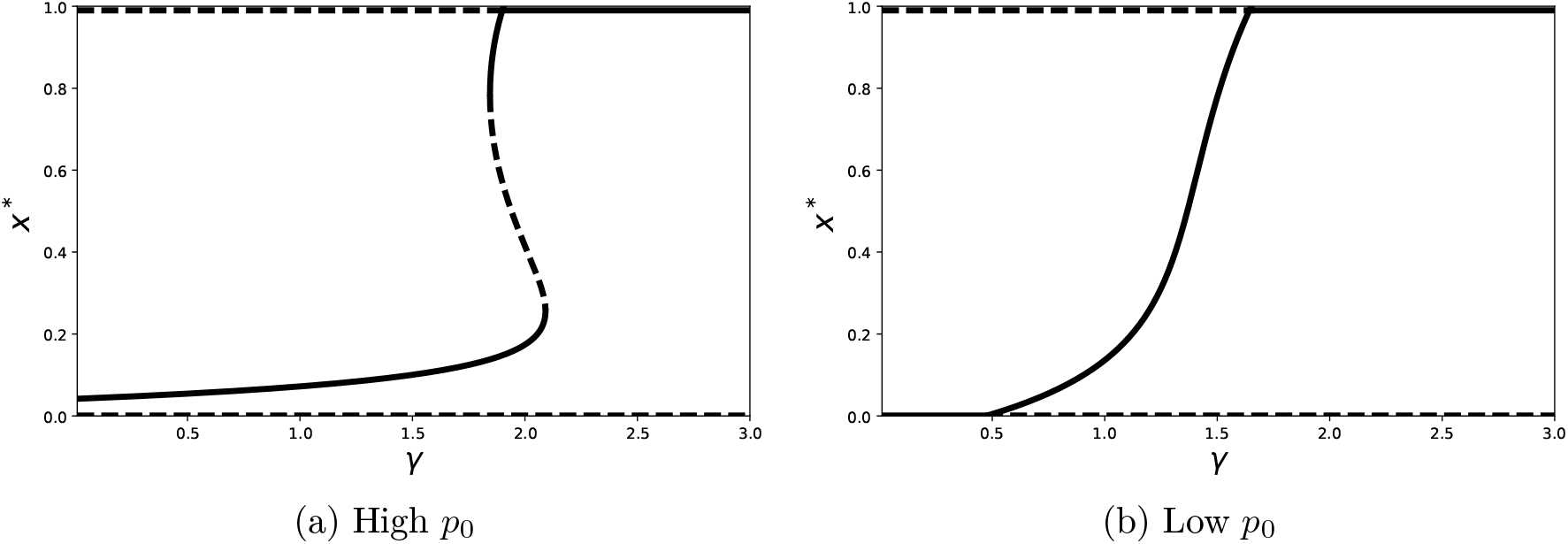
Bifurcation diagrams illustrating stable (solid line) and unstable (dashed line) equilibrium frequencies of low-effort harvesting under high and low price regimes showing multiple equilibria and hysteresis in *γ* under a high *p*_0_ and a smooth transition from high-effort dominance to low effort dominance under a low *p*_0_ regime.

**Figure S2:**
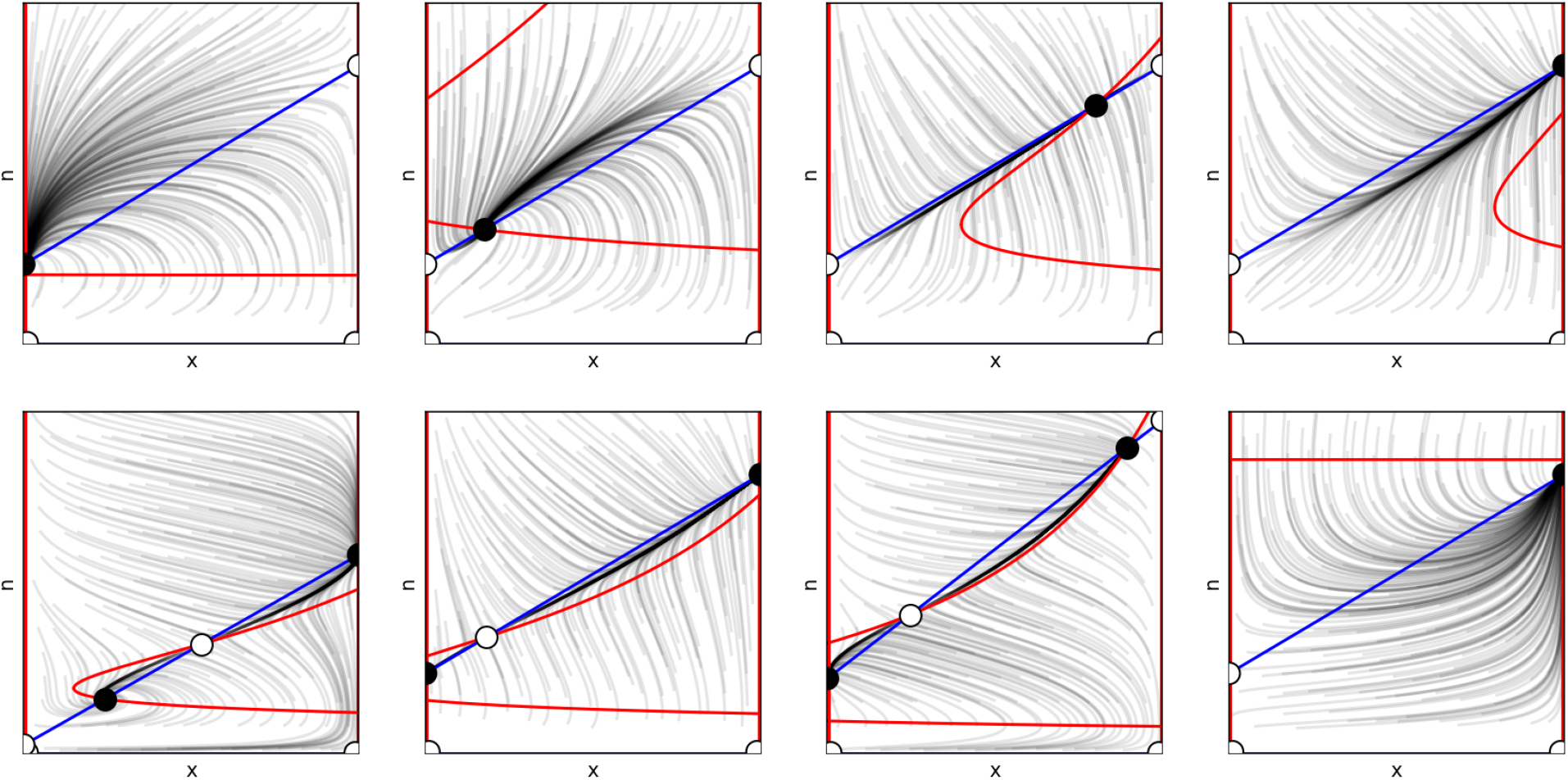
Phase planes under eight qualitatively distinct scenarios. Red curves are strategy nullclines, where 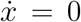. Blue lines are environment nullclines, where 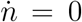. Equilibria are marked with circles, and occur at intersections of the strategy and environment nullclines. Open circles correspond to unstable equilibria, and closed circles correspond to stable equilibria.

Our model of a common-pool resource with variable harvesting efficiency is analogous to models of ostracism in common-pool systems (Tavoni et al., 2012; Tilman et al., 2017). While the details differ, both models feature added costs associated with strategies that are at low frequency – and the dynamical implications are qualitatively the same. As this analogy illustrates, there are many causes of frequency dependence in common-pool resource systems, and the framework we have developed allows the consequences to be understood generally.

Specifically, we extend the basic model of section 3.3 by incorporating frequency dependent catchability. For low effort harvesters, we let *q*_*L*_(*x*) = *q*_0_(1 + *α*_*L*_*x*) and for high effort harvesters we let *q*_*H*_ (*x*) = *q*_0_(1 + *α*_*H*_ (1 *− x*)) such that, all else being equal, harvest rates increase with increase prevalence of a strategy, and the magnitude of the benefit of increased frequency is controlled by *α*_*L*_ and *α*_*H*_. The profit of each strategy can be written as

**Figure S3:**
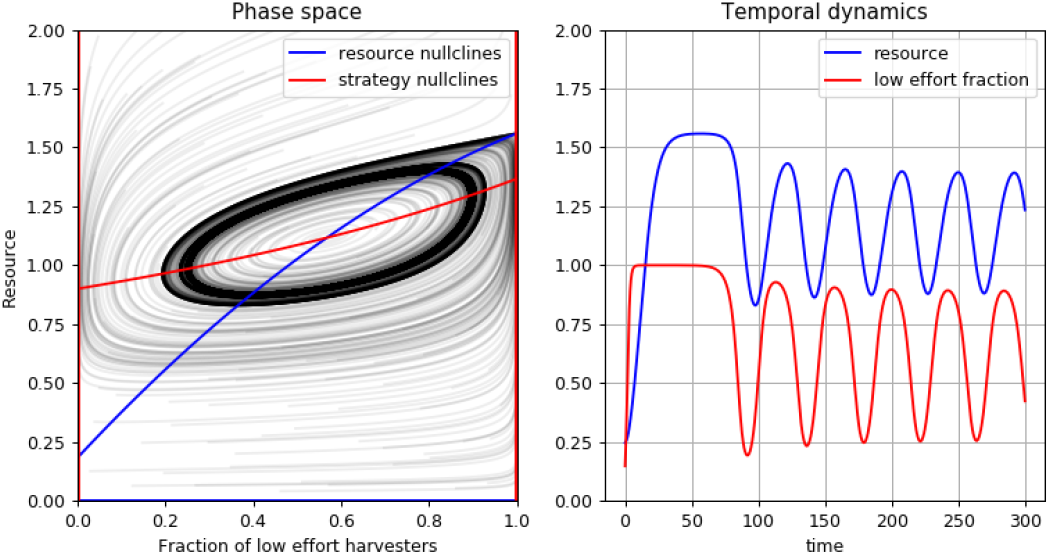
Phase plane and temporal dynamics of common-pool resource harvesting with frequency-dependent harvesting efficiency. The dynamics show convergence to a stable limit cycle. (*r* = 0.33*, K* = 4*, q* = 0.5*, e_L_* = 0.33*, e_H_* = 0.6*, p* = 10*, w* = 5*, E* = 1.3*, α*_*L*_ = 0.22, *α*_*H*_ = 0.05)

**Table 1:** Parameter values used to create each phase plane shown in Figure S1 and Figure S2, from top-left (a) to bottom-right (h)

| parameter | S1(a) | S1(b) | S2(a) | S2(b) | S2(c) | S2(d) | S2(e) | S2(f) | S2(g) | S2(h) |
| --- | --- | --- | --- | --- | --- | --- | --- | --- | --- | --- |
| $r$ | 0.3 | 0.3 | 0.3 | 0.3 | 0.3 | 0.3 | 0.36 | 0.3 | 0.36 | 0.3 |
| $k$ | 6 | 6 | 4 | 4 | 4 | 4 | 39.6 | 4 | 39.6 | 4 |
| $q$ | 0.5 | 0.5 | 0.5 | 0.5 | 0.5 | 0.5 | 0.59 | 0.5 | 0.59 | 0.5 |
| $e_L$ | 0.276 | 0.253 | 0.25 | 0.25 | 0.25 | 0.25 | 0.357 | 0.25 | 0.185 | 0.25 |
| $e_H$ | 0.564 | 0.49 | 0.5 | 0.5 | 0.5 | 0.5 | 0.6 | 0.5 | 0.515 | 0.5 |
| $p_0$ | 50 | 25 | 35 | 35 | 40 | 35 | 13.7 | 35 | 13.7 | 35 |
| $w$ | 12 | 12 | 10 | 12 | 10 | 10 | 21.7 | 5 | 16.6 | 43 |
| $\epsilon$ | 20 | 20 | 20 | 20 | 20 | 20 | 20 | 20 | 20 | 20 |
| $\gamma$ | 0-3 | 0-3 | 0.1 | 1.3 | 2.5 | 2.9 | 0.28 | 3.2 | 0.28 | 0.001 |

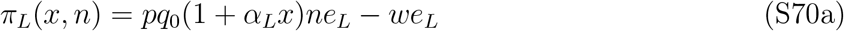

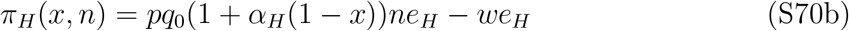

With these modifications, our system can be written as

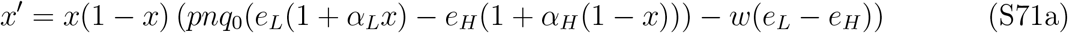

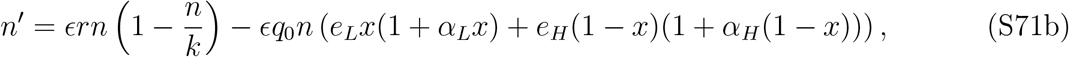

given that time has been re-scaled.

To analyze this system we consider the nullclines. The sets of points where *n*′ = 0 are *n* = 0 and

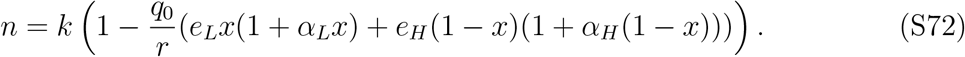

The sets of points where *x*′ = 0 are *x* = 0, *x* = 1 and

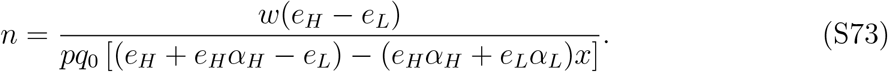

Since *e*_*H*_ > *e*_*L*_ we know that the strategy nullcline is increasing in *x* when the strategy nullcline lies within the state space, *n >* 0. Further, the resource nullcline is quadratic in *x* and opens downward. We are in particular interested in the stability of interior equilibria. Stability at interior equilibria depends on the relative timescale of resource and strategy dynamics, as well as on the slopes of the nullclines. Consider the path derivatives of the resource and strategy dynamics equations along their respective nullclines, which are equal to zero, by construction. This allows us to write the Jacobian Matrix as

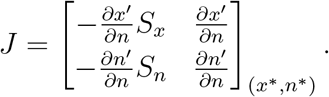

Stability at the equilibrium can be determined from the trace and determinant of *J*. We have

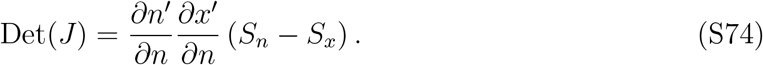

The structure of our system guarantees that 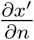 and 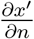 will be negative at equilibrium. Therefore, *S*_*n*_ > *S*_*x*_ is necessary, but not sufficient for stability. If *S*_*x*_ > *S*_*n*_ the interior equilibrium will be a saddle. Given that *S*_*n*_ > *S*_*x*_, then stability will occur when when the trace of *J* is negative. Due to our separation of timescales, we have a term, ∊, that controls the relative speed of resource dynamics. We have

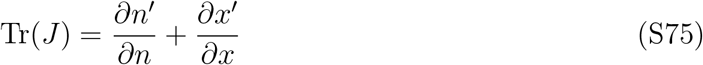

with 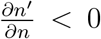 and 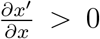. Since the first term contains ∊, we know that the trace will be negative, and stability will result for sufficiently fast resource dynamics (large ∊) and conversely the equilibrium will be unstable for slow resource dynamics.

While we can solve for the critical value of ∊ that separates parameter regions leading to stability with those leading to an unstable equilibrium and a limit-cycle, the closed form solution is unwieldy and does not enhance understanding. However, we can use this expression to quickly find the critical value of ∊ for any set of parameter values.

Changes in *r*, the intrinsic rate of growth of the resource, not only change the speed of resource dynamics, but also the location and stability of the equilibria of the system. For low *r*, corresponding to resources with low productivity, only low effort harvesting is stable. As *r* increases high effort harvesters can be supported at equilibrium, but this equilibrium is in close proximity to the low effort dominated equilibrium which is no longer stable, but only just so. Intuitively, this makes limit cycles less likely to occur because dynamics pass near two equilibria, slowing strategy dynamics and limiting environmental overshoot that drives cyclic dynamics. As a result, limit cycles occur for larger values of ∊ when the interior equilibrium is near the center of the state space. This highlights the non-monotonicity of the critical value of ∊ in *r*. For a fixed value of ∊, increasing *r* can destabilize, then stabilize the system (see Figure S4).

As *α*_*L*_ and *α*_*H*_ get larger, the strength of positive frequency dependence increases. This can lead to sets of nullclines that cross twice in the interior of the state space. At the intersection where the resource nullcline has a greater slope than the strategy nullcline stability depends on *E*, as before. The other interior equilibrium will be a saddle, and one of the edge equilibria will be stable. This can lead to three outcomes. First, if both the interior and edge equilibrium are stable, then there will be dependence on initial conditions with dynamics either leading to the interior or edge equilibrium. Next, if ∊ is small, the interior equilibrium becomes unstable and a limit cycle forms around it due to a Hopf bifurcation. In this case there is still dependence on initial conditions where either a limit cycle or an edge equilibrium result. Finally, if ∊ is too small, the interior equilibrium is unstable and no limit cycle results, all initial conditions lead to the edge equilibrium.

To illustrate this complexity, we consider a range of cases with where only *r* changes. When we let ∊ = 2.1, increasing *r* can first de-stabilize the interior equilibrium, then stabilize it. This highlights the non-monotonicity of the critical value of ∊ in *r*. Figure S4 shows that intermediate values of *r* can lead to cyclic dynamics. This is in contrast to the stability condition under the renewable resource model where we found that both *r* and ∊ had similar effects of stability.

**Figure S4:**
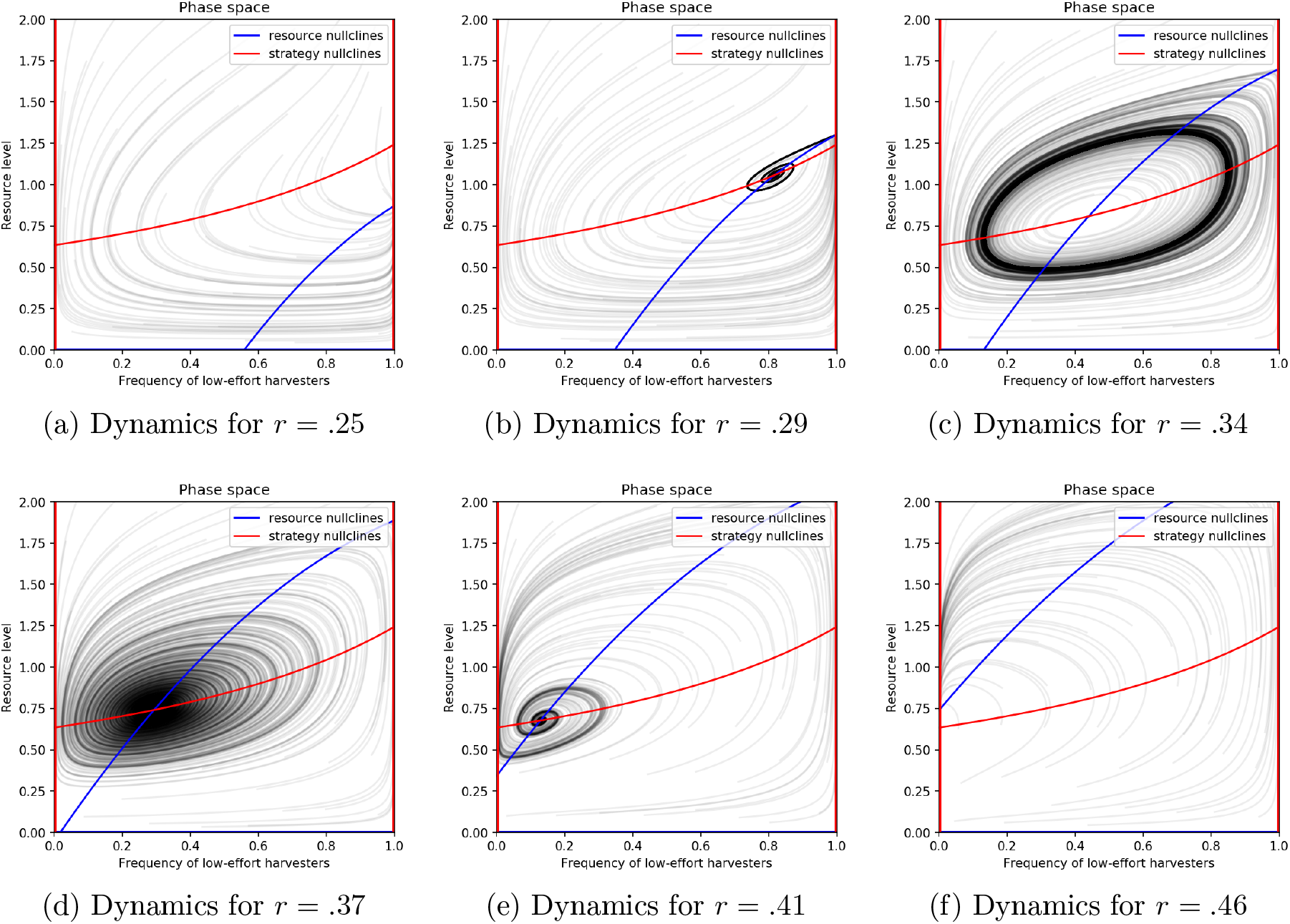
As *r*, the intrinsic rate of growth of the resource, increases, the speed of resource dynamics increase and the location of the equilibria change. As opposed to changes in ∊ where higher values always stabilize an interior equilibrium, increasing *r* can either stabilize or de-stabilize an interior equilibrium depending on context. Here we show that all else being equal, increase *r* can lead from stable equilibria, to cycles and back. (∊ =. 25*, q* =. 5*, K* = 4, *p* = 10, *w* = 5, *e*_*L*_ =. 34, *e*_*H*_ =. 6, *α*_*L*_ =. 15, *α*_*H*_ =. 25)

